# Designing a *Saccharomyces cerevisiae* Cell Factory for the Overaccumulation of Tyrosol using Rational Design and Genome-Scale Metabolic Modeling Approaches

**DOI:** 10.64898/2026.11.23.768537

**Authors:** Alba Guillén, Fabián Moreno-Avitia, Sandra Sánchez-Martí, Ricardo Bisquert, José M. Guillamón

## Abstract

Tyrosol (TyOH) is a phenolic compound with antioxidant and anti-inflammatory properties, naturally present in two Mediterranean products, wine and virgin olive oil, and is the key precursor of Hydroxytyrosol or Salidroside. Its industrial production still relies on chemical synthesis, which conflicts with the EU Green Deal roadmap to transition to a clean, circular, resource-efficient economy by 2050. Thus, moving to a sustainable economy is a priority. Here, we engineered *Saccharomyces cerevisiae* as a cell factory to produce TyOH. To support this goal, we used ecFactory, a pipeline leveraging enzyme-constrained genome-scale metabolic models (ecGEMs), to predict gene targets, narrowing the candidate list and simplifying experimental validation for strain development. The modifications were tested in a medium composed mainly of glucose with ammonium sulfate as the sole nitrogen source. The final strain reached 876.91 ± 20.37 mg·L^-1^ in a 0.5 L bioreactor, demonstrating the potential of model-guided engineering for sustainable TyOH production.

## 1. Introduction

Aromatic amino acids (AAA), including tyrosine (Tyr), phenylalanine (Phe), and tryptophan (Trp), are synthesized through the shikimate pathway and subsequently catabolized by yeasts, as in other eukaryotes, via the Ehrlich pathway (Maeda & Dudareva, 2012; Mas et al., 2014). During alcoholic fermentation (AF), this route releases AAA-derived fusel alcohols as part of nitrogen metabolism, including 2-phenylethanol (2PE), tryptophol, and TyOH, the latter being the final higher alcohol formed from Tyr (Cordente et al., 2021; Hazelwood et al., 2008). Consequently, these pathways release bioactive compounds of significant commercial interest (Rumpl et al., 2025), making them attractive targets for biotechnological production (Li et al., 2025).

TyOH can be found in extra-virgin olive oil (EVOO), where it arises due to the natural hydrolysis of secoiridoids present in fresh olives, such as oleuropein and ligstroside, which together account for 90 % of the initial polyphenol content (Karkoula et al., 2014; López de las Hazas et al., 2016). TyOH and its derivatives, above all hydroxytyrosol (HT), exhibit potent antioxidant and anti-inflammatory properties (Robles-Almazan et al., 2018), contributing to the health benefits associated with the Mediterranean diet, which are characterized by regular intake of olive oil and its derivatives (Daniele et al., 2017). Accordingly, the European Food Safety Authority (EFSA) approved a health claim in 2011, later incorporated into European Regulation (EU) No. 432/2012 (European Commission, 2012), supporting the consumption of EVOO containing at least 5 mg of HT and its derivatives per 20 g of oil. TyOH has also been reported to exhibit antimicrobial, neuroprotective, and cardioprotective properties (Plotnikov & Plotnikova, 2020; Shen & Mao, 2026), supporting its use as a functional ingredient in nutraceuticals, cosmetics, and pharmaceutical formulations (Karković Marković et al., 2019). More broadly, the market for Tyr-derived compounds is estimated to be in the billion-dollar range, reflecting their broad applications across the food, cosmetics, and pharmaceutical industries (Guo et al., 2023).

Despite this growing interest, TyOH production remains mainly performed via chemical synthesis from petroleum-derived precursors or by extraction from natural sources such as olive mill wastewater and olive oil by-products, relying on both low yields and poor recovery rates (Junker et al., 2025; Sannino et al., 2013; Zahi et al., 2022). Both strategies present significant drawbacks: chemical synthesis typically involves hazardous reagents, multiple purification steps, higher costs, and a substantial environmental footprint, while extraction from plant matrices is constrained by low natural abundance, seasonal availability, and batch-to-batch variability (Shen et al., 2024). These methods also require large amounts of starting biomass and release harmful intermediates, such as acetyl gluco-oligosaccharides and acetylsalidroside (Britton et al., 2019; C. Li et al., 2026). Consequently, these limitations have prompted a shift towards bio-based production using engineered microbial cell factories (MCFs), which enable TyOH biosynthesis under controlled, reproducible conditions with lower environmental impact (Bisquert et al., 2022).

As a Generally Recognized as Safe (GRAS) microorganism with a well-characterized genome, extensively studied metabolism, and a broad genetic toolbox (Long et al., 2024; Nielsen, 2019), *Saccharomyces cerevisiae* (*S. cerevisiae*) offers a highly tractable, robust, and versatile platform for strain engineering. This versatility has been exploited across diverse industries, ranging from traditional food and beverage fermentation sectors to more recent biotechnological applications as a sustainable alternative for the production of biofuels, pharmaceuticals, nutraceuticals, or chemicals, among others (Cunha et al., 2020; Hong & Nielsen, 2012; Walker & Stewart, 2016). In this context, *S. cerevisiae* stands out as a particularly suitable host for TyOH production (Borodina & Nielsen, 2014; Guo et al., 2019). Nevertheless, the synthesis of AAA-derived compounds is not a primary metabolic function of yeast and key enzymes are subject to feedback inhibition (Luttik et al., 2008), so efficient production requires metabolic engineering of the strain and optimization of the cultivation process (Liu et al., 2019; Rahmat & Kang, 2020).

Rational engineering of the Ehrlich route requires extensive prior biochemical knowledge to identify the steps and enzymes to modulate, and is limited by our incomplete understanding of cellular regulation and strain behavior. Systems biology offers a complementary approach. In *S. cerevisiae,* genome-scale metabolic models (GEMs) have been widely used to guide strain optimization for the production of diverse value-added compounds, including aromatic metabolites (Gold et al., 2015; J. Zhang et al., 2020). By integrating known reactions, metabolites, and gene–protein–reaction associations, GEMs predict flux distributions under defined constraints and identify bottlenecks and deletion and overexpression targets, thereby narrowing the experimental search space before strain construction (He & Lu, 2025; Zhang & Hua, 2016). Their predictive power, however, depends on the completeness of the reconstruction, and previous studies have identified gaps in the representation of TyOH biosynthesis in yeast models (Henriques et al., 2021; Scott et al., 2020). The community-consensus model Yeast9 integrates accumulated biochemical and physiological knowledge (Zhang et al., 2024), while enzyme-constrained GEMs (ecGEMs), such as ecYeastGEM, further incorporate enzyme catalytic capacities and proteome allocation constraints, providing a more physiologically informed view of flux limitations (Chen et al., 2024a; Lu et al., 2019).

Building on ecYeastGEM and the GECKO toolbox, the ecFactory pipeline integrates enzyme-constrained modeling with computational target selection. It yields a reduced list of gene targets associated with enhanced production, accounting for each enzyme’s position in the synthetic flux and the competing demands of growth, without disrupting essential cellular functions. The framework has been applied to predict targets for 103 valuable compounds, including tyrosol (TyOH), although these predictions have not been systematically validated experimentally (Domenzain et al., 2025). Such strategies can reduce the experimental effort required to develop yeast cell factories for aromatic compounds (Ishchuka et al., 2022). Model predictions are, however, limited to the information encoded in the model, and rational design based on prior knowledge can complement them, maximizing the resulting designs.

Here, we addressed the lack of experimental validation by combining ecFactory predictions with rationally selected modifications, building on our previous experience engineering yeast for other aromatic bioactive compounds, to develop a *S. cerevisiae* BY4743 strain for the de novo production of TyOH from glucose. The 13 selected targets, located in the shikimate and Ehrlich pathways and in competing routes, were sequentially integrated via overexpression, down-regulation, or deletion. Each strain was characterized for TyOH production and growth in small-scale cultures before scale-up to a 0.5 L bioreactor, and the experimental data were used to assess the agreement between model predictions and observed results.

## 2. Materials and Methods

**2.1. Strains and Culture Conditions:**

The *S. cerevisiae* strains used in the present study are described in **Table 1**. The laboratory strain BY4743, provided by Euroscarf, was used as a chassis strain for successive genetic modifications to improve TyOH titers. Yeast strains were grown and cultivated using YPD medium [20 g·L^-1^ glucose, 20 g·L^-1^ peptone, 10 g·L^-1^ yeast extract], in SC medium [20 g·L^-1^ glucose, 5 g·L^-1^ ammonium sulphate, 1.7 g·L^-1^ Yeast Nitrogen Base (YNB) without amino acids and ammonium sulphate (Difco), and an indicated amount of a supplement drop-out powder (Formedium)] or in SD medium [20 g·L^-1^ glucose, 5 g·L^-1^ ammonium sulphate, 1.7 g·L^-1^ Yeast Nitrogen Base (YNB) without amino acids and ammonium sulphate (Difco)]. All media were supplemented with 18 g·L-1 agar (NZYTech, Lisbon, Portugal) to prepare solid media at 28°C. To test the effect of modifications aimed at over-accumulating TyOH, the selected medium for the shake-flask production experiments was SD medium with 80 g·L^-1^ glucose (SD80). Depending on each strain’s auxotrophic needs (**Table 1**), media were supplemented with histidine (20 mg·L^-1^), uracil (20 mg·L^-1^), or leucine (120 mg·L^-1^). For overnight cultures, cells were grown at 150 rpm and 28°C.

**Table 1.**
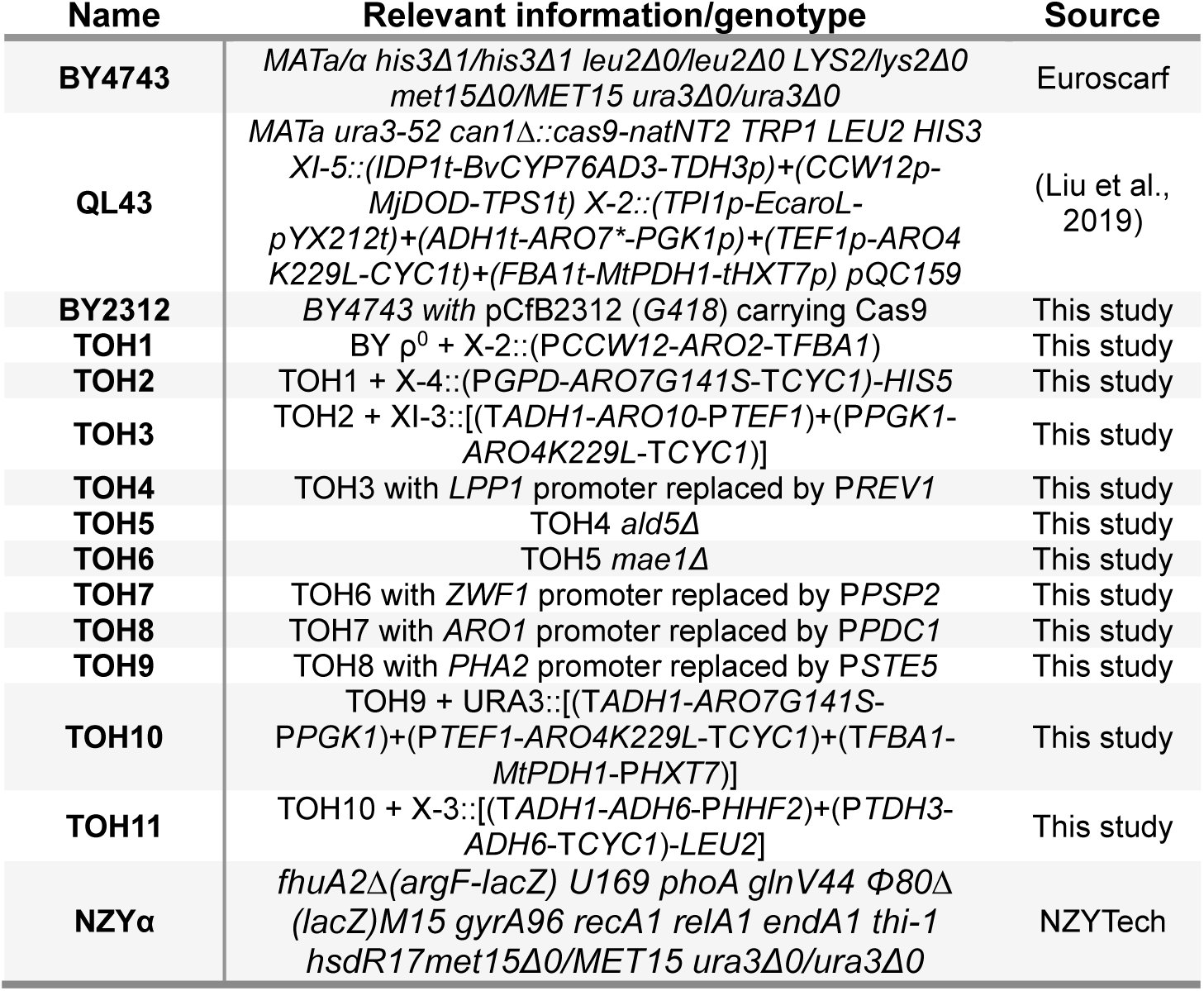
List of yeast strains used in this work.

| Name | Relevant information/genotype | Source |
|---|---|---|
| **BY4743** | *MATa/α his3Δ1/his3Δ1 leu2Δ0/leu2Δ0 LYS2/lys2Δ0 met15Δ0/MET15 ura3Δ0/ura3Δ0* | Euroscarf |
| **QL43** | *MATa ura3-52 can1Δ::cas9-natNT2 TRP1 LEU2 HIS3 XI-5::(IDP1t-BvCYP76AD3-TDH3p)+(CCW12p-MjDOD-TPS1t) X-2::(TPI1p-EcaroL-pYX212t)+(ADH1t-ARO7\*-PGK1p)+(TEF1p-ARO4 K229L-CYC1t)+(FBA1t-MtPDH1-tHXT7p) pQC159* | (Liu et al., 2019) |
| **BY2312** | *BY4743 with* pCfB2312 (*G418*) carrying Cas9 | This study |
| **TOH1** | BY ρ<sup>0</sup> + X-2::(P*CCW12-ARO2*-T*FBA1*) | This study |
| **TOH2** | TOH1 + X-4::(P*GPD-ARO7G141S*-T*CYC1)-HIS5* | This study |
| **TOH3** | TOH2 + XI-3::[(T*ADH1-ARO10*-P*TEF1*)+(P*PGK1-ARO4K229L*-T*CYC1*)] | This study |
| **TOH4** | TOH3 with *LPP1* promoter replaced by P*REV1* | This study |
| **TOH5** | TOH4 *ald5Δ* | This study |
| **TOH6** | TOH5 *mae1Δ* | This study |
| **TOH7** | TOH6 with *ZWF1* promoter replaced by P*PSP2* | This study |
| **TOH8** | TOH7 with *ARO1* promoter replaced by P*PDC1* | This study |
| **TOH9** | TOH8 with *PHA2* promoter replaced by P*STE5* | This study |
| **TOH10** | TOH9 + URA3::[(T*ADH1-ARO7G141S*-P*PGK1*)+(P*TEF1-ARO4K229L*-T*CYC1*)+(T*FBA1-MtPDH1*-PHXT7)] | This study |
| **TOH11** | TOH10 + X-3::[(T*ADH1-ADH6*-P*HHF2*)+(P*TDH3-ADH6*-T*CYC1*)-*LEU2*] | This study |
| **NZYα** | *fhuA2Δ(argF-lacZ) U169 phoA glnV44 Φ80Δ(lacZ)M15 gyrA96 recA1 relA1 endA1 thi-1 hsdR17met15Δ0/MET15 ura3Δ0/ura3Δ0* | NZYTech |

**Table 2.**
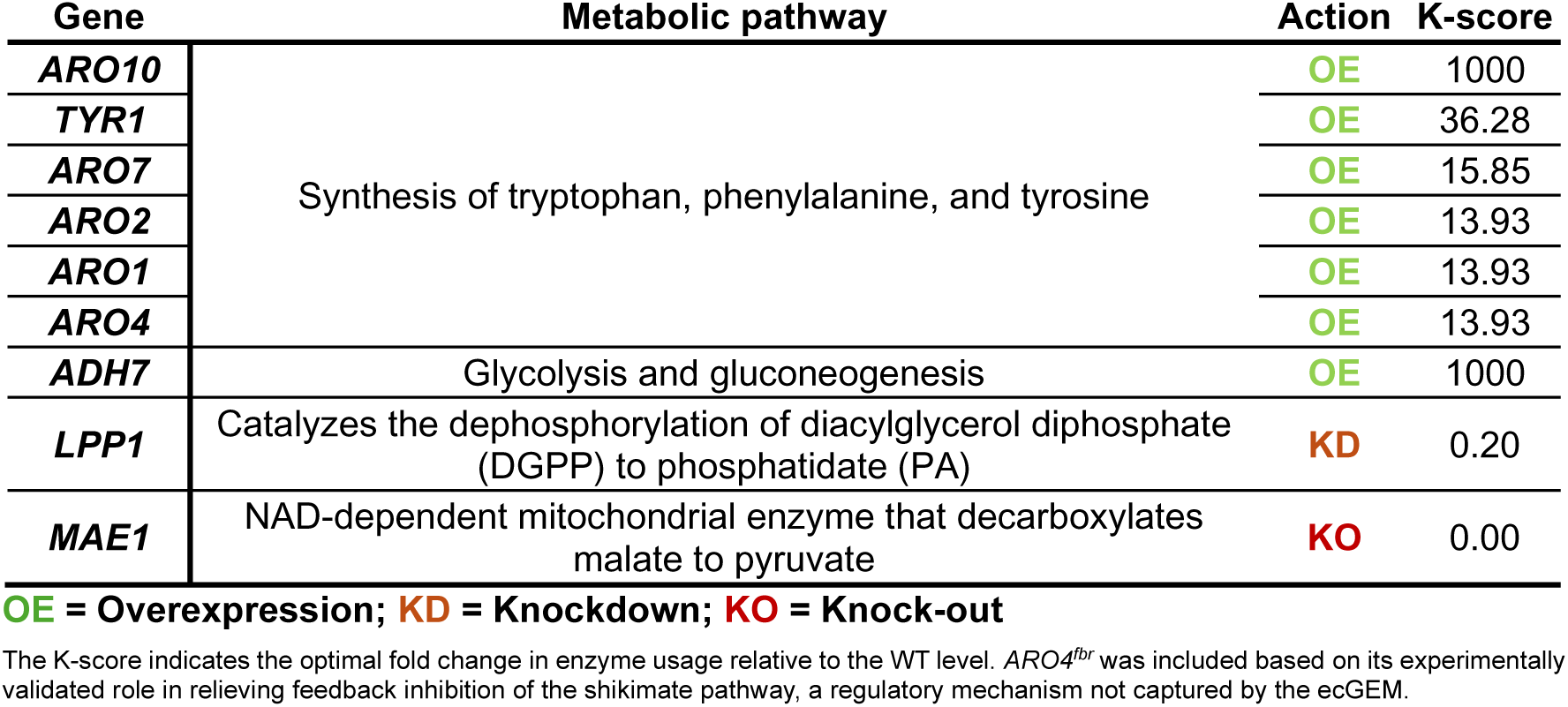
ecFactory-predicted engineering targets for TyOH production.

| **Gene** | **Metabolic pathway** | **Action** | **K-score** |
|---|---|---|---|
| ***ARO10*** | Synthesis of tryptophan, phenylalanine, and tyrosine | **OE** | 1000 |
| ***TYR1*** |  | **OE** | 36.28 |
| ***ARO7*** |  | **OE** | 15.85 |
| ***ARO2*** |  | **OE** | 13.93 |
| ***ARO1*** |  | **OE** | 13.93 |
| ***ARO4*** |  | **OE** | 13.93 |
| ***ADH7*** | Glycolysis and gluconeogenesis | **OE** | 1000 |
| ***LPP1*** | Catalyzes the dephosphorylation of diacylglycerol diphosphate (DGPP) to phosphatidate (PA) | **KD** | 0.20 |
| ***MAE1*** | NAD-dependent mitochondrial enzyme that decarboxylates malate to pyruvate | **KO** | 0.00 |

**OE = Overexpression; KD = Knockdown; KO = Knock-out**
The K-score indicates the optimal fold change in enzyme usage relative to the WT level. *ARO4<sup>fbr</sup>* was included based on its experimentally validated role in relieving feedback inhibition of the shikimate pathway, a regulatory mechanism not captured by the ecGEM.

As a cloning host for plasmid construction and maintenance, the strain *Escherichia coli* (*E. coli*) NZYα (NZYTech, Lisbon, Portugal) was used, being cultured in lysogeny broth (LB) medium [10 g·L^-1^ tryptone, 5 g·L^-1^ yeast extract and 5 g·L^-1^ sodium chloride (NaCl)] supplemented with 100 mg·L^-1^ ampicillin or 50 mg·L^-1^ kanamycin to maintain plasmids at 37°C in a labroller at 400 rpm.

### 2.2. Constructed Strains and Plasmids

All the genetically modified strains derived from BY4743 are listed in **Table 1**. The strains constitute a cumulative series (TOH1 to TOH11), in which each strain carries all modifications of the preceding one plus one additional modification, introduced in the order shown in **Table 3**. All the plasmids constructed and primers employed in this study are listed in **Tables S1 and S2**, respectively. The genomic DNA of the yeast strain BY4743 was extracted according to the DNA extraction method described by Querol et al. (1992), and all the endogenous genes and regulatory elements (including promoters and terminators) were amplified from the genomic DNA of our parental strain or from the MoClo toolkit plasmid collection (Lee et al., 2015). Genes up to 4 kb were amplified using Phusion High-Fidelity DNA polymerase (Thermo Scientific), while DNA fragments bigger than 4 kb were amplified using 2 x Phanta Flash Master Mix (Vazyme Biotech, China). DNA from agarose gels or PCR products was purified using the NZYGelpure kit (NZYTech, Lisbon, Portugal), and DNA quantity was quantified using a NanoDrop.

**Table 3.** Stepwise genetic modifications implemented during TyOH-strain construction.

| **Blocks** | **Strain** | **Applied modification** | **Accumulated genotype** |
|---|---|---|---|
| **-** | **BY4743** | pCfB2312 | none |
| **B1** | **TOH1** | OE *ARO2* | WT +*ARO2* |
|  | **TOH2** | OE *ARO7<sup>G141S</sup>* | TOH1 + *ARO7*\* |
|  | **TOH3** | OE (*ARO10-ARO4<sup>K229L</sup>*) | TOH2 + *ARO10* + *ARO4*\* |
| **B2** | **TOH4** | KD *LPP1* | TOH3 + *lpp1*-KD |
|  | **TOH5** | KO *ALD5* | TOH4 + *ald5Δ* |
|  | **TOH6** | KO *MAE1* | TOH5+ *mae1Δ* |
|  | **TOH7** | KD *ZWF1* | TOH6 + *zwf1*-KD |
|  | **TOH8** | OE *ARO1* | TOH7 + *ARO1* |
| **B3** | **TOH9** | KD *PHA2* | TOH8 + *pha2*-KD |
|  | **TOH10** | OE (*ADH6-ADH7*) | TOH9 + *ADH6* + *ADH7* |
|  | **TOH11** | OE *MtPDH1* | TOH10 + *MtPDH1* |

#### 2.2.1. Design and Construction of Repair Fragments

Gene knock-outs (KO) and knock-downs (KD) were generated by CRISPR-Cas9-mediated genome editing using a linear repair donor. Repair fragments for KO were made by designing a forward primer, including 50 nucleotides upstream of ATG + 10 nucleotides downstream the STOP codon of the gene, and a reverse complement primer consistent in 50 nucleotides downstream STOP + 10 nucleotides upstream ATG. Both primers had a total length of 60 nucleotides. The donor fragment was amplified using a high-fidelity polymerase via partially overlapping primers, also known as “template-free PCR”. For KD, the native promoter was replaced with a weak constitutive promoter from the MoClo toolkit of Lee et al. (2015) for modulating the activity of the selected genes. Donors were amplified using a forward primer including 40 nucleotides upstream of the ATG of the native promoter region + 20 nucleotides from the start of the new promoter, and a reverse primer consistent with 40 nucleotides upstream of the ATG of the gene of interest + 20 nucleotides upstream of the end of the new promoter. All donors were amplified with Phusion High-Fidelity DNA Polymerase (Thermo Scientific) and purified from the PCR products prior to transformation.

#### 2.2.2. Construction of Overexpression (OE) Cassettes and Cloning into Site-directed Integrative Vectors

Gene cassettes designed for OE were cloned into integrative vector backbones from the EasyClone collection (Jensen et al., 2014) and assembled into the EasyClone-MarkerFree toolkit (Jessop-Fabre et al., 2016), which directs CRISPR-Cas9-mediated integration at defined genomic loci without selection markers (**Table S2**). Otherwise, to restore the auxotrophies present in our strain, the single-copy integrative vectors pCfB257 and pCfB258 carrying the *Kluyveromyces lactis* leucine (*KlLEU2*) and *Schizosaccharomyces pombe* histidine (*SpHIS5*) markers, respectively, were also used (**Table S2**).

In order to perform the cloning of the ARO genes from *S. cerevisiae*, *ARO2* was amplified from the genomic DNA of BY4743, while *ARO10* and the feedback inhibition-insensitive *ARO4^K229L^*(*ARO4*\*) and *ARO7^G141S^* (*ARO7*\*) were amplified from a previously constructed p423GPDp vector (Bisquert et al., 2022). The designed primers (**Table S2**) for amplifying the particular PCR biobricks contained a single uracil base and were amplified using Phusion U Hot Start polymerase (Thermo Scientific). This uracil tail allows the assembly via Uracil-Specific Excision Reaction (USER) cloning technology. In parallel, the selected EasyClone vectors were prepared by sequential treatment with FastDigest enzyme *Asi*SI (*Sfa*AI) (Thermo Fisher Scientific) and a nicking endonuclease *Bsm*I (New England Biolabs). After purification, the USER-ready vector and the PCR biobricks were mixed at a molar ratio of 1:3 (vector:insert) and treated with the USER^TM^ enzyme mix (New England Biolabs). Additionally, to avoid using the same terminators and reduce the likelihood of homologous recombination, some of the OE cassettes were constructed via overlapping PCR (**Table S2**) to amplify promoters, genes, terminators, and integrative sequences with overlapping ends.

All ARO genes were overexpressed under strong constitutive promoters, where *ARO10* and *ARO4\** were cloned into pCfB2904, *ARO7*\* was cloned into pCfB258, while *ARO2* was designed via fusion PCR. On the other hand, to fine-tune the flux of precursors towards TyOH, the *ADH6* and *ADH7* dehydrogenases were amplified from the yeast genome and later cloned into the pCfB257 plasmid and the prephenate dehydrogenase *PDH1* from *Medicago truncatula* (Mt*PDH1*), an analogue of *S. cerevisiae TYR1,* was amplified from strain XJ020 (Jiao et al., 2024) and cloned into the integrative vector pYTK096 from the MOCLO toolkit, thereby complementing the auxotrophies of the strain, due to the presence of the URA3 marker.

After the cloning was complete, the mixture was used directly for bacterial transformation. Successful cloning of our constructs was confirmed by colony PCR in *E. coli* isolates using the recommended *ADH1*_test-F and *CYC1*_test-R primers for final constructs smaller than 5 kb, while different primer pairs were used for larger constructs (**Table S2**). To verify the exact sequence of the insert, Sanger sequencing (Eurofins Genomics, Ebersberg, Germany) was performed. Prior to yeast transformation, the resulting vectors were linearized with FastDigest *Not*I (Thermo Fisher Scientific) and subsequently purified from agarose gels.

#### 2.2.3. Design and Assembly of Cas9-sgRNA Plasmids

For CRISPR-Cas9 targeting, the target spacer was designed using the CHOPCHOP web tool (Labun et al., 2019). All single-guide RNA (sgRNA) sequences created are listed in Table S3. The sgRNAs were ligated by Golden Gate assembly using type IIS restriction enzymes into the pWS series vectors from the Ellis laboratory (available at Addgene: https://www.addgene.org/browse/article/25503/). These Cas9 gap repair vectors are designed to facilitate sgRNA cloning. Briefly, complementary oligonucleotides encoding the desired sgRNA target sequence were annealed into the pWS backbone, bearing the Cas9 expression cassette and the sgRNA scaffold. The oligonucleotide mixture is incubated in a one-tube reaction with the plasmid, the enzyme *Bsm*BI (NEB), and the T4 DNA ligase (Thermo). The ligation product is transformed into NZY5α *E. coli* cells. The successful ligations were identified by the absence of fluorescence, due to the substitution of sfGFP at the cloning site, allowing rapid identification of successful sgRNA insertions. This strategy allows efficient generation of customized CRISPR plasmids.

### 2.3. Yeast Transformation

Once the expression cassettes were constructed and purified, either to integrate, downregulate, or delete, the resulting linear DNA donors were transformed into the yeast genome following the PEG/Lithium Acetate (LiAc) method (Gietz & Woods, 2002). Briefly, 150 μL of overnight-saturated yeast cultures were inoculated into 5 mL of fresh SC selection medium and incubated at 28°C until the total biomass reached 2.5 × 10^8^ cells. Harvest the cells at 4000 g at room temperature for 5 min, wash the cells in 1 mL of sterile water into 1.5 mL microcentrifuge tubes, and then centrifuge at 13000 rpm for 30 s. After discarding the supernatant, cells were washed with LiAc (0.1 M) and centrifuged at 13000 rpm for 1 min. Cells were pelleted and resuspended with 240 μL of PEG 3350 (50% w/v), 36 μL of LiAc (1.0 M), 50 μL of single-stranded carrier DNA (2.0 mg/mL), and 34 μL of sterile water plus the linearized donor. The parental strain BY4743 was first transformed with plasmid pCfB2312 for constitutive Cas9 expression (BY2312). For each genetic modification, approximately 1.5–2 µg of the corresponding linearized fragment from the integrative vector or KD/KO donors was used for yeast transformation, whereas 400–500 ng of the resulting Cas9/sgRNA vectors were used. Yeast cells were selected on selective agar medium based on strain-specific auxotrophic markers and plasmid maintenance requirements. The subsequent transformants were checked by colony PCR to determine whether the modifications were performed correctly in both alleles, in one allele, or not at all. Once the colonies were confirmed, these strains were cryostocked in SC medium with 35% glycerol.

### 2.4. Cultivations

#### 2.4.1. Growth Conditions for Tyrosol Quantification

To test the effect of the different modifications performed in our engineered strain and their influence on Tyr accumulation. Precultures of each modified strain were grown overnight at 28°C with orbital shaking at 150 rpm in 5 mL of SD culture media supplemented with the required amino acid to complement auxotrophic markers, in case it had not yet been covered. Then the culture was further inoculated into 100 mL glass shake flasks containing 30 mL of fresh SD80 production medium, with an initial OD_600_ of 0.2. The incubation was set up at 28°C with orbital shaking at 150 rpm and was grown for 120 h. All the experiments were carried out in biological triplicates. Final OD_600_ was measured and recorded using a spectrophotometer (Cell Density Meter model 40, Fisher Scientific) after diluting the sample to a range of 0.2–0.7. Then TyOH levels were determined by collecting 10 mL of supernatant by centrifugation at 4000 x g for 5 min at room temperature in a benchtop 5804-R centrifuge with a rotor for plates (Eppendorf, Leipzig, Germany).

#### 2.4.2. Bioreactor Cultivations

To verify the robustness of the final modified strain, two sets of fermentations were carried out. In this way, we tested the strain’s growth relative to its parent, measuring glucose and fermentation by-product levels, and then determining the final concentration of TyOH produced by HPLC. Batch cultures were grown at 28°C in three 0.5 L reactors (MiniBio, Applikon Biotechnology) with a working volume of 0.25 L of SD80 medium. A temperature probe connected to a Peltier cryostat controlled the temperature of the cultures. The stirrer was set to 350 rpm, and the pH was set to 5.0 and controlled with 3 M KOH. Each bioreactor was initially inoculated with 1·10^6^ cells·mL^-1^ of the parental strain and the definitive TyOH overproducer strain (TOH11). Precultures were grown overnight in SC medium at 28 °C. Biomass accumulation was monitored using OD_600_ to track the process and calculate the maximum growth rate (μ_max_). In addition, cell dry weight (CDW) was measured in triplicate at several time points throughout the cultivation. A 5 mL aliquot of each sample was collected to measure CDW, and the samples were then filtered through a 0.45 µm membrane filter. Afterward, filters were dried for 20 min at maximum power in a microwave oven and then weighed on an analytical balance (ABS220-4N, Kern). Samples were collected for further analysis by removing the pellet, and the supernatant was stored at -20 °C.

### 2.5. Determination of Aromatic Higher Alcohols and Tyrosol

#### 2.5.1. Determination of Extracellular Tyrosol Concentration by HPLC-FLUO

A volume from the fermentations was transferred to 12 mL tubes, and the supernatant was collected by removing the cells after centrifugation at 4000 × g for 5 minutes at room temperature. Then, samples were diluted 50% v/v with HPLC-grade absolute methanol, mixed by vortexing, and centrifuged for 1 min at 13000 rpm. The supernatant was filtered through a 0.22 µm nylon filter before the chromatographic analysis. Tyr was analyzed by high-performance liquid chromatography (HPLC) on a Waters ACQ Arc Sys Core chromatograph (Waters, Milford, MA, USA) coupled with a Waters 2475 fluorescence detector (FLUO), a quaternary pump, an autosampler and an online degasser. Chromatographic separation was carried out in an Accuore^TM^ C18 (Thermo Scientific, Waltham, Massachusetts, USA) (4.6 × 150 mm and a 2.6 μm particle size) column. The mobile phases used were A (0.1 % formic acid in acetonitrile) and B (0.1 % formic acid in ultrapure water), with a constant flow rate of 1 mL/min. The injection volume was 10 µL, and the gradient program was as follows: 0 – 15 min, 100 % B (0 % A), 15 – 16 min 90 % B (10 % B), 16 – 17 min 0 % B (100 % A) 17 – 20min 100 % B (0 % A). The column temperature was set at 30°C and samples were held at 12°C. The FLUO detector was set at λ_ex_ = 280 nm and at λ_em_ = 320 nm. The retention time for Tyr with this configuration was 10.216 min.

#### 2.5.2. Determination of Extracellular Higher Alcohols by Gas Chromatography (CG)

Higher alcohols and esters, including 2-phenylethanol (2PE), were analyzed using a TRACE 1600 gas chromatograph fitted with a flame ionization detector (FID) and a TriPlus RSH autosampler (Thermo Scientific, Waltham, MA, USA). Volatile compounds were extracted via the headspace solid-phase microextraction (HS-SPME) technique using a 100-µm polydimethylsiloxane (PDMS) fiber (Supelco, Sigma-Aldrich, Spain). The carrier gas was helium (1 mL/min), and the GC column was an Agilent HP INNOWax 30 m × 0.25 m capillary column coated with a 0.25-m layer of cross-linked polyethylene glycol (Agilent Technologies). The CG analysis was performed according to the method previously described by Minebois et al. (2025), and the instrumental settings and parameters were applied as reported herein. For sample preparation, 5 mL of supernatant was transferred into a 20 mL vial containing 5 mL of NaCl saline solution (75 g NaCl in 247.5 mL ultrapure water), and the vial was tightly closed with a PTFE septum plug.

### 2.6. Bioinformatic and Statistical analysis

#### 2.6.1. Calculation of specific growth rate values

All the constructed TyOH overproducer strains were tested in a 48-well microtiter plate (Nunc^TM^, Thermo Fisher) and their growth was measured using a SPECTROstar Nano® microplate reader (BGM Labtech, Offenburg, Germany) at 28 °C, measured at a 600 nm wavelength with 15 s of 400 rpm double-orbital shaking before each reading. The optical density was measured every 30 min until they reached the stationary phase. Each well was filled with 700 μL of SD80 medium to complement the required auxotrophies and inoculated with an initial OD_600_ of 0.1 from a previously saturated overnight preculture. In addition, for bioreactors batch cultures, the OD_600_ was taken during all the process with the same objective. The OD values obtained over time were used to calculate the kinetic parameters of each strain following the Gompertz equation (Zwietering et al., 1990):

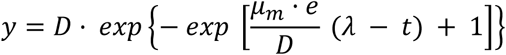

The μ_max_, the maximal population (D) and the lag phase period (λ) were obtained using R software (4.5.0), while the area under the growth curve (AUC) was determined for all the different strains using Origin Pro 2019b.

#### 2.6.2. Modeling the tyrosol flux using the ecFactory pipeline

The TyOH production model was derived from the batch version of the enzyme-constrained *S. cerevisiae* model, ecYeastGEM (Domenzain et al., 2025). Model editing and simulations were performed in MATLAB R2025a (version 25.1.0) using the model-editing functions of RAVEN Toolbox (v2.11.0) (Wang et al., 2018). All analyses described here were carried out in a WT context strain. TyOH biosynthesis was incorporated into the base model via the native yeast Ehrlich pathway. The reactions already present in ecYeastGEM convert Tyr to 4-hydroxyphenylpyruvate (4-HPP) through native reactions. Two reactions were added to complete product formation: (i) decarboxylation of 4-HPP to 4-hydroxyphenylacetaldehyde (4-HPAA) by the phenylpyruvate decarboxylase *ARO10* (YDR380W), and (ii) reduction of 4-HPAA to TyOH by the NADPH-dependent aldehyde reductase *ADH7* (YCR105W). The ecFactory code, input models, and complete results, including the final candidate list (**Table S3**), are available on GitHub (https://github.com/fma91/Tyrosol_ecYeast).

#### 2.6.3. Prediction and prioritization of engineered targets

Engineering targets were predicted with ecFactory using the GECKO 2.0.3 implementation of enzyme-constrained flux scanning with enforced objective function (ecFSEOF), followed by enzyme-usage variability analysis (EUVA). All optimization problems were solved in MATLAB R2025a using the RAVEN Toolbox 2 with the Gurobi Optimizer 12.0.3 (Chen et al., 2024).

The ecYeast model was constrained with medium and glucose as the carbon source. ecFSEOF was run over 16 enforced-biomass steps. The scanning interval extended from 0.5 to 2 times the experimental yield. Genes with K-scores of at least 1.05 were classified as overexpression candidates, genes with K-scores below 0.5 as downregulation candidates, and genes with K-scores below 0.05 as knockout candidates. Targets with intermediate K-scores were removed. Essential genes were excluded from the KO set.

The remaining candidates were classified using a variability analysis. For the production state, glucose uptake was fixed at 1 mmol·g_DCW_^-1^·h^-1^, biomass formation was fixed to the yield specified above, and TyOH production was constrained to its maximum attainable value. Minimum, maximum, and parsimonious enzyme usages were then calculated for each candidate. A reference state was generated by requiring TyOH production to remain above 1 % of its maximum and by maximizing biomass formation at the same glucose uptake rate. Candidates with inconsistent changes between the production and reference states were removed. The remaining targets were prioritized based on the distance between their enzyme-demand ranges. Finally, combinations of targets were tested mechanistically while retaining at least 99 % of the prescribed biomass flux, and a minimal target combination was selected. These successive stages produced the level-1, level-2 and level-3 candidate tables. Transport reactions predicted by ecFSEOF but lacking enzyme associations were reported separately.

#### 2.6.4. Reconstruction of the cumulative strain series and production-envelopes analysis

To emulate iterative strain construction in the laboratory, a cumulative genotype series (TOH) was defined *in silico*, in which each successive strain carries all edits of the preceding strain plus one additional ecFactory-prioritized modification (**Table 3**). The cumulative modifications were the OE of *ARO2* in TOH1; additional *ARO7* in TOH2; additional *ARO4* and *ARO10* in TOH3; *LPP1* KD in TOH4; *ALD5* KO in TOH5; *MAE1* KO in TOH6; *ZWF1* KD in TOH7; *ARO1* OE in TOH8; *PHA2* KD in TOH9; and *ADH6* and *ADH7* OE in TOH10. Finally, TOH11 was derived from TOH10 by OE of *MtPDH1*.

*In silico* genetic modifications were implemented by changing enzyme-usage bounds. Basal usage was obtained from the maximum enzyme usage (maxUsageBio) in the biomass-reference state of the level-2 (L2) candidate list, and was increased by 1 % to avoid numerical equality with the optimum. For the product-envelope analysis, OE was represented by a lower enzyme-usage bound equal to the basal usage multiplied by the specified factor either derived from the ecFactory scan (approximately 14-fold for *ARO2*, *ARO7*, *ARO4*, and *ARO1*) or a fixed factor for *ARO10*, *ADH6*, *ADH7*, and *TYR1*. KD was represented by an upper bound of 21 % of basal usage. If an edit prevented growth at unit glucose uptake, the knockdown cap was relaxed to the enzyme usage required at the growth optimum. KO edits removed the corresponding gene from the model.

#### 2.6.5. Dynamic FBA of the definitive TOH11 batch culture

Batch dynamics of the final strain, TOH11, were simulated using dynamic flux balance analysis (dFBA) with the ecYeastGEM model and the complete cumulative genotype. The simulation was implemented in MATLAB R2025a using RAVEN Toolbox 2, GECKO 2.0.3, and Gurobi Optimizer 12.0.3. OE increased enzyme capacity without imposing a minimum usage flux, whereas the KD bounds were multiplied by a global scaling factor calibrated to the measured growth rate. Initial glucose and biomass concentrations were obtained from the experimental time course. Biomass formation was maximized at each time step. TyOH production was subsequently maximized in a second optimization while maintaining at least 90 % of the maximum growth rate.

## 3. Results and discussions

### 3.1. Enzyme-constrained model-guided strain design for tyrosol production

In recent years, we have focused on developing efficient yeast platforms for the overproduction of bioactive compounds, particularly those related to the aromatic amino acids Trp and Tyr (Bisquert et al., 2022; Planells-Cárcel et al., 2025). Predictive tools based on ecGEM models assist in creating consistent MCFs using well-established, up-to-date yeast models. The ecFactory framework was applied to the ecGEM to identify gene engineering targets for TyOH overproduction by comparing enzyme usage between the biomass-optimal and TyOH-production-optimal solutions across the phenotypic phase plane. Candidate targets were ranked across three stringency levels (L1–L3), and the final modification set for the definitive strain (TOH11) was assembled by selecting L3 targets (highest priority), supplementing with biologically motivated L2 candidates, and excluding targets with overlapping or redundant roles in ecYeastGEM (Table S3).

Five overexpression targets were identified as essential and tightly constrained across all production conditions (L3). Four encode consecutive enzymes of the shikimate and aromatic amino acid biosynthesis pathway: *ARO2* (chorismate synthase, OE k-score 13.9), *ARO7* (chorismate mutase, k-score 15.9), *ARO1* (pentafunctional AROM polypeptide, k-score 13.9), and *TYR1* (prephenate dehydrogenase, k-score 36.3). Their classification reflects that tyrosol production requires substantially greater aromatic precursor flux than biomass synthesis alone can sustain. The fifth target is the Ehrlich pathway enzyme *ARO10* (4-hydroxyphenylpyruvate decarboxylase), and the final committed step, *ADH7* (4-hydroxyphenylacetaldehyde reductase), which together convert 4-hydroxyphenylpyruvate to TyOH. One knockdown target was predicted: *LPP1* (lipid phosphate phosphatase, KD k-score 0.21), whose enzyme usage is coupled to biomass and competes with the shared protein pool under production conditions. *MAE1* (mitochondrial malate dehydrogenase) was predicted to be a knockout and classified as unnecessary for production.

*ARO4* (DAHP synthase, Tyr-sensitive isoform) appeared as a supporting overexpression target at the L2 set but did not reach the L3 target gene set because it is not essential for TyOH production *in silico*. Despite *ARO4*’s ∼9.5-fold higher enzyme usage during TyOH production, which would theoretically indicate higher yield, its usage dropped to zero at some points in the production envelope. This indicates that the *ARO3* isoform can substitute for the activity of the other isoform. This result, combined with the high *k_cat_* of *ARO4* (122 s⁻¹), renders its increased enzyme demand non-essential during the simulations. Thus, *ARO4* is beneficial but not rate-limiting. However, feedback inhibition is a variable not yet incorporated into ecModels. Therefore, the need for or the impact of a feedback-resistant variant of *ARO4* (*ARO4*^fbr^) cannot be predicted by our approach, although its overexpression has been shown to positively impact TyOH production. *ARO4^fbr^*was incorporated in the definitive modification target set due to its reported positive impact on aromatic compound production by maintaining active carbon flux through the shikimate pathway, even during the accumulation of aromatic amino acids (Rodriguez et al., 2015).

#### 3.1.1. Overexpression (OE) targets identified with the ecFactory model

The selection of *ARO4* and *ARO7* is consistent with previous rational metabolic engineering efforts reported in *S. cerevisiae* and with experimental data showing that both enzymes represent the two major regulatory bottlenecks of the pathway (Luttik et al., 2008). Aro4p is a DAHP synthase (3-Deoxy-D-arabinoheptulonate 7-phosphate), which catalyzes the first committed step of chorismate biosynthesis, the aldol condensation of phosphoenolpyruvate (PEP) and erythrose-4-phosphate (E4P), and is feedback-inhibited by Tyr. The Tyr branch-point enzyme Aro7p, a chorismate mutase, is likewise inhibited by Tyr and activated by Trp. Overexpression (OE) of feedback-insensitive variants has been shown to increase the production of compounds derived from the Ehrlich pathway by diverting more carbon to this aromatic pathway, specifically raising the TyOH titers approximately 3-fold relative to wild-type alleles (Cordente et al., 2019; Koopman et al., 2012; Liu et al., 2019).

The remaining downstream of *ARO4* genes, *ARO1* and *ARO2*, catalyze several steps in the shikimate pathway and encode the pentafunctional AROM enzyme and chorismate synthase, respectively (Duncan et al., 1987; Gottardi et al., 2017). These two enzymes act sequentially between DAHP and chorismate, and their selection as OE targets reflects the need to reinforce flux throughout the entire shikimate pathway rather than at a single rate-limiting node, ensuring that the increased carbon entry enabled by *ARO4* deregulation is efficiently funneled toward chorismate (CHA) without generating new intermediate bottlenecks. Immediately downstream of the shikimate pathway is *TYR1,* a NAD(P)^+^ dependent prephenate dehydrogenase that catalyzes the committed step converting prephenate (PPA) into 4-hydroxyphenylpyruvate (4-HPP), the last dedicated step before transamination to Tyr (Mao et al., 2017). Its selection as an OE target is aimed at reinforcing flux specifically toward Tyr, rather than the other AAA, which is in line with the reported strategies for producing TyOH, where *TYR1*, together with feedback-resistant *ARO4* and *ARO7,* improves the yield, underscoring *TYR1* as a key node for coupling upstream aromatic precursor supply (Guo et al., 2019). Nevertheless, the native *TYR1* gene product is subject to feedback inhibition by Tyr, as is *ARO4*. Consequently, the OE of this enzyme would be expected to have a limited impact on flux toward TyOH, as *Tyr1p* activity would remain constrained by the end-product amount. To overcome this limitation, *TYR1* OE was instead performed using the *Medicago truncatula PDH1* ortholog (*MtPDH1*), a cytosolic prephenate dehydrogenase reported to be insensitive to Tyr-mediated inhibition, thereby alleviating this possible limitation in the metabolic flux through TyOH (Liu et al., 2019; Schenck et al., 2020).

With the flux strongly redirected toward the Tyr branch, the model predicted the overexpression of *ARO10* and *ADH7*. Notably, these two genes were the only directly pathway-associated enzymes recovered in the top-priority gene set, as they function in the final two steps of the Ehrlich pathway, converting the 4-HPP derived from Tyr into hydroxyphenylacetaldehyde (4-HPAA) by *ARO10* and then *ADH7,* catalyzing the reduction of 4-HPAA to TyOH (Gottardi et al., 2017). This result was consistent with the underlying model architecture, which restricts TyOH synthesis to the native Ehrlich route (Guo et al., 2019). Within this route, *ARO10* encodes the decarboxylase, which catalyzes the first divergently specific step to the conversion of Tyr towards TyOH (Hazelwood et al., 2008). The inclusion of *ADH7* in the model appears to reflect its significant ability to efficiently convert 4-HPAA to TyOH, as *ADH7*, together with *ADH6* and *ADH4*, has been reported to show the highest substrate specificity for this conversion step (Li et al., 2018; Ruan et al., 2022). However, they are not the most canonical and highly expressed ADH isoforms in yeast central metabolism, but have been specifically implicated in the reduction of aromatic and branched-chain aldehydes (Larroy et al., 2002; Yang et al., 2018). Together, the joint selection of *ARO10* and *ADH7* as top up-regulation targets, alongside the reinforcement of the entire upstream shikimate-to-Tyr route, reflects a coherent push-pull strategy in which upstream precursor supply is increased in parallel with the improved capacity to convert those precursors into the final target compound.

#### 3.1.2. Downregulation and knock-out (KD/KO) targets identified by ecFactory

The top downregulation/knockout targets (*MAE1*, *LPP1*) corresponded to reactions that compete for carbon or redox flux outside the TyOH pathway. *MAE1* is a non-essential gene that encodes a NAD-dependent mitochondrial malic enzyme, which catalyzes the oxidative decarboxylation of malate to pyruvate. *MAE1* has been proposed to contribute to intramitochondrial NADPH or pyruvate provision in anaerobic metabolism. However, null mutants showed that this function is dispensable under aerobic conditions (Boles et al., 1998). Its knock-out (KO) has previously been identified, together with other TCA-cycle and respiration-associated genes, as part of a strategy to reduce TCA cycle and respiratory fluxes, allocating carbon away from acetyl-CoA and derived fatty acid synthesis, amino acids derived from 2-oxoglutarate or nucleotide biosynthesis, in favor of several flavonoid production (Domenzain et al., 2025). Thus, its selection by the ecFactory pipeline is supported as an optimal target that redirects flux away from mitochondrial respiration toward cytosolic aromatic amino acid biosynthesis.

*LPP1* is a membrane protein related to glycerolipid metabolism, which encodes a Mg^2+^-independent lipid phosphate phosphatase that catalyzes the dephosphorylation of bioactive lipid intermediates such as phosphatidate (PA) or lyso-PA (LPA) that yields diacylglycerol (DAG), an intermediate for the synthesis of triacylglycerols (Furneisen & Carman, 2000; Toke et al., 1998). Furthermore, *LPP1* may be involved in lipid signaling and has been reported to hydrolyze farnesyl diphosphate (FPP) to farnesol, suggesting it could be a potential engineering target for the synthesis of isoprenoids and terpenoids in yeast (Faulkner et al., 1999; Wang et al., 2011). In yeast metabolic engineering, *LPP1*, together with *DPP1*, has proven effective for overaccumulating isoprenoid-derived products, such as geranylgeraniol, via OE (Tokuhiro et al., 2009). In contrast, the KO of *LPP1* has been used as a strategy to prevent the dephosphorylation and consequent loss of key terpenoid precursors, redirecting them toward heterologous prenyltransferases and terpene synthases, an approach that has been successfully applied to increase the titers of structurally complex terpenoids such as (+)-borneol or ophiobolins (Zhang et al., 2024; Zhang et al., 2025).

Unlike *MAE1*, the functional link of *LPP1* to aromatic amino acid or TyOH biosynthesis is not well documented in the literature, and its identification as a KD target likely reflects the combinatorial, model-driven nature of the ecFactory search rather than a previously established rational design principle. Nevertheless, a plausible mechanistic explanation can be proposed, given that after the KD of *LPP1,* a reduction in the flux towards the branch of the reserve lipid synthesis (triacylglycerols) would be expected. This could conserve acyl-CoA and the associated NADPH/ATP consumption normally directed to the biosynthesis of storage lipids, releasing cofactors and carbon for other biosynthetic requirements, in line with the general pattern observed among the other downregulated targets identified by the model.

### 3.2. *Rationally engineering the L-Tyr branch to avoid competitive routes for* tyrosol biosynthesis

In parallel to the ecFactory-derived targets, four additional genetic modifications (*ADH6, ALD5, PHA2*, and *ZWF1*) were proposed based on rational design principles and literature evidence, prior to the implementation of the ecGEM-based workflow (**Table 2**).

To achieve the maximum conversion of 4-HPAA to TyOH, the OE of *ADH6*, along with the *ADH7* isoform identified by the model, was proposed. Given the competence that exists for 4-HPAA as a substrate, which can also natively be converted to 4-hydroxyphenylacetate (4-HPA) by the action of any aldehyde reductase (ALD), losing part of the increased pool of suppliers, the introduction of a new ADH could help us to enhance the final step of the TyOH synthesis. *ADH6* was chosen among the other isoforms based on its high catalytic efficiency toward aromatic and branched-chain aldehydes, including 4-HPAA, as demonstrated by Ruan et al. (2022). In this framework, the model’s preference for *ADH7* over *ADH6*, despite both being NADPH-dependent enzymes of nearly identical size (*ADH6*, UniProt Q04894, 360 aa; *ADH7*, 361 aa), likely reflects differences in the isoform-specific ***k_cat_*** values employed by the model for both enzymes based, rather than differences in cofactor usage or protein size/synthesis cost. Under the GECKO/ecModel formulation, in which enzyme usage is constrained by the ratio between flux and ***k_cat_*** together with a limited total protein pool (Domenzain et al., 2022; Sánchez et al., 2017), even nearly identical, cofactor-equivalent isoforms can be differentially prioritized by the model based on relatively small differences in catalytic efficiency toward the specific substrate considered.

As part of our efforts to improve the TyOH titer in our yeast, we aimed to modulate as many competitive pathways for key substrates as possible, thereby preventing potential leakage without compromising the predictions of our pipeline. As we mentioned previously, 4-HPAA can be converted to its derived acid by several endogenous ALDs. However, any *S. cerevisiae* ALD has been unambiguously characterized as performing this specific reaction on 4-HPAA, so modulating this competitive pathway was not an easy task given this lack of annotation. In contrast, *E. coli* possesses a well-characterized phenylacetaldehyde dehydrogenase, *feaB* (P80668), which catalyzes this competing oxidation to form 4-HPA. Taking this into account, we performed a structural homology-based search (using the DALI server) to identify a *feaB* paralog in yeast, finding that *ALD5* showed the highest structural similarity to feaB, followed by *ALD4*, which has also been reported elsewhere as a potential candidate for the oxidation of different phenolic aldehydes (Datta et al., 2017). Beyond this structural argument, functional evidence further supports the involvement of Ald5p in the catabolism of phenolic aldehydes, as *ALD5*, together with *PAD1*, *ATF1,* and *ATF2*, has been shown to participate in the conversion of phenolic aldehydes and acids such as coniferyl aldehyde, ferulic acid and p-coumaric acid in *S. cerevisiae*, with TyOH identified among the most abundant products detected in the corresponding strains, supporting a broader role for Ald5p in phenolic aldehyde metabolism beyond its originally described function in mitochondrial acetate formation and ethanol consumption after fermentation (Adeboye et al., 2017). Considering this, the *ALD5* allele was KO to preserve as much 4-HPAA as possible, avoiding the loss of this substrate, which is necessary for its conversion to TyOH. The Ehrlich pathway also leads to the synthesis of other aromatic higher alcohols derived from Phe, such as 2-phenylethanol (2PE). This compound shares its precursor node with Tyr, as both amino acids derive from PPA, the branch-point metabolite downstream CHA (Braus, 1991). *PHA2*, encoding prephenate dehydratase, was targeted to eliminate competing flux toward the Phe branch at the diverging node, redirecting carbon flux toward Tyr and, subsequently, through TyOH biosynthesis. This same strategy has been applied in *S. cerevisiae* to boost Tyr-derived products such as hydroxytyrosol (HT) in our previous research (Bisquert et al., 2022). This modification, however, must be carefully balanced against the risk of generating a Phe auxotroph. Then, a KD was chosen as the best option to avoid the funneling of the enhanced PPA pool without needing to supplement the media with an extra amino acid, compromising the cost and scalability of the production in the future.

Finally, in order to optimize the synthesis of E4P, *ZWF1* encoding glucose-6-phosphate dehydrogenase was selected for KD to reduce flux through the oxidative branch of the PPP branch, thereby promoting a more carbon-conserving distribution of flux toward E4P and PEP, the two precursors required to enter the shikimate pathway, where E4P is typically the limiting metabolite (Becker & Boles, 2003; Gottardi et al., 2017). This strategy has been previously implemented with the OE of *TKL1* for rewiring the PPP to reverse flux from the glycolytic intermediates fructose-6-phosphate (F6P) and glycerol-3-phosphate (G3P) to E4P and xylulose-5-phosphate (X5P), relieving constraints on flux entering the AAA biosynthetic route by up to seven-fold (Deaner & Alper, 2017). Furthermore, its deletion has been associated with lower fatty acid production, reflecting the broader metabolic cost of reduced NADPH availability beyond the aromatic pathway itself (Cadière et al., 2011). Despite this, the KD or KO of *ZWF1* directly compromises the oxidative PPP, the main cellular source of NADPH, leading to NADPH deficiency and forcing the cell to couple its growth to Tyr production via the OE of the NADP^+^-dependent *TYR1* allele (Gold et al., 2015). However, this cofactor deficiency can become problematic under conditions of oxidative stress or redox imbalance, and for this reason, its net benefit is not always guaranteed. Taken together, these model-derived and rationally designed targets provided the complete set of genetic tools that were gradually integrated into the TyOH production strain, as described in the following section.

### 3.3. *Metabolic engineering of the TyOH strain: implementation and product* characterization

A total of 13 metabolic targets, derived from both ecFactory predictions and rational design based on our results and the previous literature, were selected to engineer our laboratory strain for the *de novo* production of TyOH from sole carbon sources. These modifications were integrated sequentially, yielding the final cumulative series TOH1 to TOH11 (genotypes in **Table 1**). The final strain, TOH11, increased TyOH production by almost 80-fold relative to the parental strain BY4743. All strains were cultured in parallel under identical shake-flask conditions, enabling direct comparison of TyOH titer, sugar consumption, and biomass accumulation across the series. Because the modifications were not introduced in the order in which they are best understood, their individual and cumulative effects are presented in three blocks grouped by metabolic rationale, in line with the framework introduced above, rather than following the chronology of construction (**Table 3**).

**Figure 1** summarizes how these targets relate to the shikimate and Ehrlich pathways and to Tyr-derivative synthesis, as well as to the competing routes that divert carbon, redox or cofactor flux away from TyOH biosynthesis.

**Figure 1.**
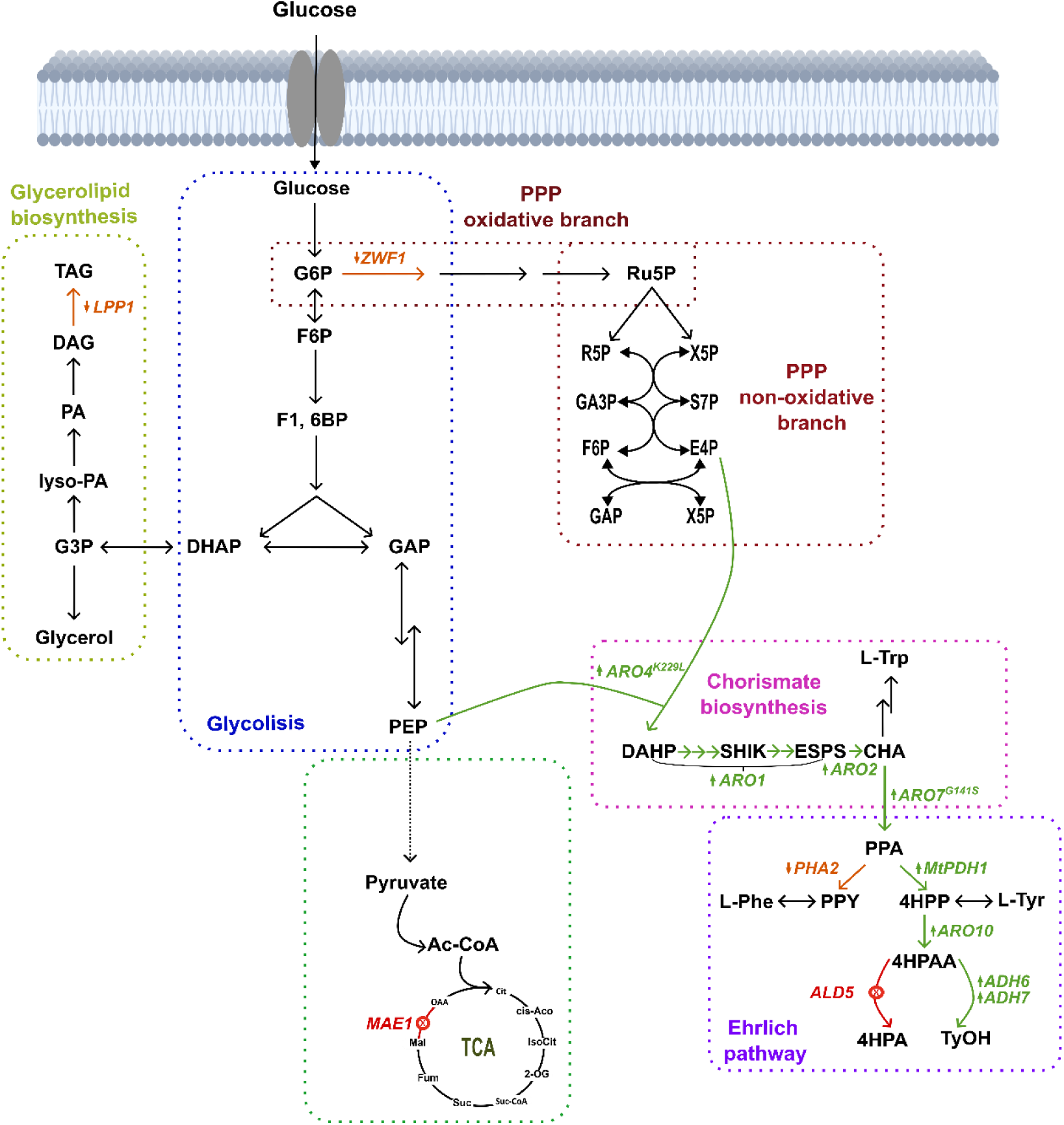
Overview of engineered yeast central metabolism linking glucose uptake to central carbon metabolism to enhance TyOH biosynthesis. Central carbon metabolism connects glycolysis where glucose is sequentially converted to pyruvate through key intermediates such as G6P glucose-6-phosphate, F6P fructose-6-phosphate, F1,6BP fructose 1,6-biphosphate, DHAP dihydroxyacetone phosphate, GAP glyceraldehyde 3-phosphate and PEP phosphoenolpyruvate; and the pentose phosphate pathway (PPP), comprising its oxidative branch, where G6P is oxidized to ribulose-5-phosphate (Ru5P) with concomitant NADPH generation, and its non-oxidative branch, which regenerates glycolytic intermediates such as R5P ribose-5-phosphate, X5P xylulose-5-phosphate, GA3P glyceraldehyde 3-phosphate, S7P sedoheptulose 7-phosphate, G3P glyceraldehyde 3-phosphate, and X5P xylulose 5-phosphate, as well as, supplies erythrose-4-phosphate (E4P). The condensation of E4P and PEP initiates the shikimate (SHIK) pathway, proceeding through key intermediates including 3-deoxy-D-arabino-2-heptulosonic acid 7-phosphate (DAHP) and chorismate (CHA), the branch-point metabolite leading to aromatic amino acid biosynthesis, Trp L-tryptophan, Phe L-phenylalanine, and Tyr L-tyrosine, culminating in higher alcohols production via the Ehrlich pathway. The TCA cycle involves cycling through Cit citrate, cis-Aco cis-aconitate, IsoCit isocitrate, 2-OG 2-oxoglutarate, Suc-CoA succinyl-CoA, Suc succinate, Fum fumarate, Mal malate, and OAA oxaloacetate, which serves as the primary hub for energy generation and the supply of biosynthetic precursors. The triacylglycerol biosynthesis pathway is shown branching from G3P toward the synthesis of fatty acids and membrane phospholipids, a highly ATP-and NADPH-demanding anabolic route whose flux is modulated by LPP1 through lipid phosphate dephosphorylation. Overexpressed yeast endogenous genes are shown in green, including Tyr-feedback-insensitive DAHP synthase *(ARO4*^K229L^), pentafunctional aromatic protein (*ARO1*), chorismate synthase (*ARO2*), Tyr-feedback-insensitive chorismate mutase *(ARO7*^G141S^), phenylpyruvate decarboxylase (*ARO10*), and alcohol dehydrogenases (*ADH6* and *ADH7*). In addition, the heterologous L-tyrosine prephenate dehydrogenase-encoding gene (Mt*PDH1*) from Medicago truncatula was evaluated for enhancing L-tyrosine supply. Downregulated genes are shown in orange, including prephenate dehydratase (*PHA2*), glucose-6-phosphate dehydrogenase (*ZWF1*), and lipid phosphate phosphatase (*LPP1*). Removed endogenous mitochondrial genes are shown in red, including aldehyde dehydrogenase (*ALD5*) and the malic enzyme (*MAE1*). DHQ 3-dehydroquinate, DHS 3-dehydro-shikimate, SHIK shikimate, S3P shikimate-3-phosphate, EPSP 5-enolpyruvyl-shikimate-3-phosphate, PPY phenylpyruvate, 4*-*HPP 4-hydroxyphenylpyruvate, 4-HPAA 4-hydroxyphenyl acetaldehyde, 4-HPA 4-hydroxyphenylacetate.

#### 3.3.1. Push-pull engineering of the shikimate and Ehrlich pathways (B1)

The production of TyOH in yeast can only be accomplished via the Tyr (*TAL* branch) synthesis of the Ehrlich pathway. This route is known to be highly accurate. Thus, improving the expression levels of the responsible enzymes at each step can be readily achieved with appropriate genetic tools (**Figure 1**). As detailed previously, *ARO2*, *ARO7*, *ARO10,* and *ARO4* were identified by the ecFactory pipeline as top-priority up-regulation targets, jointly reinforcing carbon entry into the shikimate pathway and alleviating Tyr by-product inhibition (when insensitive alleles are used to bypass feedback inhibition), thereby enhancing pathway-specific flux toward this aromatic pathway. These four genes were the first to be introduced into the BYCas9 background, establishing the initial push-pull framework upon which subsequent modifications were built. The physiological parameters for each cultivation, along with production titers of TyOH and 2PE, are shown in **Table 4**.

**Table 4.** Product formation during shake-flask cultivations of the engineered strain series.

| **Work blocks** | **Strain** | **X (OD<sub>600</sub>)** | **X (g· L<sup>-1</sup>)** | **TyOH titer<br>(mg· L<sup>-1</sup>)** | **2PE titer<br>(mg·L<sup>-1</sup>)** |
|---|---|---|---|---|---|
| **B1-2-3** | **WT** | 5.17 ± 0.15<sup>bcd</sup> | 1.16 ± 0.03<sup>bcd</sup> | 11.56 ± 0.43<sup>a</sup> | 34.12 ± 3.20<sup>a</sup> |
| **B1** | **TOH1** | 5.43 ± 0.06<sup>d</sup> | 1.22 ± 0.01<sup>d</sup> | 33.19 ± 0.21<sup>a</sup> | 44.29 ± 4.33<sup>a</sup> |
|  | **TOH2** | 5.37 ± 0.06<sup>cd</sup> | 1.21 ± 0.01<sup>cd</sup> | 63.22 ± 1.72<sup>a</sup> | 65.05 ± 4.95<sup>a</sup> |
|  | **TOH3** | 5.47 ± 0.21<sup>d</sup> | 1.23 ± 0.05<sup>d</sup> | 273.41 ± 18.50<sup>b</sup> | 371.31 ± 1.39<sup>d</sup> |
| **B2** | **TOH4** | 5.53 ± 0.15<sup>d</sup> | 1.24 ± 0.03<sup>d</sup> | 331.83 ± 8.25<sup>c</sup> | 415.72 ± 4.77<sup>d</sup> |
|  | **TOH5** | 5.50 ± 0.26<sup>d</sup> | 1.24 ± 0.06<sup>d</sup> | 391.05 ± 15.10<sup>d</sup> | 661.53 ± 10.69<sup>f</sup> |
|  | **TOH6** | 6.13 ± 0.15<sup>e</sup> | 1.37 ± 0.03<sup>e</sup> | 347.58 ± 10.06<sup>cd</sup> | 643.56 ± 13.52<sup>f</sup> |
|  | **TOH7** | 3.63 ± 0.15<sup>a</sup> | 0.83 ± 0.03<sup>a</sup> | 328.83 ± 10.38<sup>c</sup> | 251.96 ± 18.63<sup>c</sup> |
|  | **TOH8** | 4.13 ± 0.12<sup>a</sup> | 0.94 ± 0.03<sup>a</sup> | 375.42 ± 30.66<sup>cd</sup> | 359.28 ± 5.61<sup>d</sup> |
| **B3** | **TOH9** | 4.87 ± 0.23<sup>bc</sup> | 1.10 ± 0.05<sup>bc</sup> | 555.79 ± 20.23<sup>e</sup> | 347.59 ± 42.88<sup>d</sup> |
|  | **TOH10** | 4.73 ± 0.21<sup>b</sup> | 1.07 ± 0.05<sup>b</sup> | 788.73 ± 36.84<sup>f</sup> | 181.40 ± 4.85<sup>b</sup> |
|  | **TOH11** | 6.20 ± 0.20<sup>e</sup> | 1.39 ± 0.04<sup>e</sup> | 902.97 ± 19.43<sup>g</sup> | 432.96 ± 23.10<sup>e</sup> |

Total biomass (X) was assessed by measuring OD<sub>600</sub> and CDW, whereas the amounts of TyOH and 2PE are expressed in mg·L<sup>-1</sup>. Data are presented as mean ± SD (*n* = 3 independent biological replicates per strain). Statistical differences were assessed independently for each column using one-way ANOVA followed by Tukey's post hoc test (*p* < 0.05). Within a column, values sharing the same letter are not significantly different. Letters are assigned from the lowest (a) to the highest mean value.

Overexpression of *ARO2* (strain TOH1) increased the TyOH titer from 11.56 ± 0.44 to 33.19 ± 0.21 mg·L^-1^ (**Table 4**), representing a 2.87-fold increase relative to the BY4743 wild-type strain, consistent with the de-bottlenecking of the final step in CHA biosynthesis from 3-enolpyruvyl-shikimate 5-phosphate (ESPS), allowing the entry of more carbon to the Ehrlich pathway. Subsequent introduction of the feedback-resistant *ARO7* (*ARO7\**) allele (strain TOH2) resulted in an increase of 5.15-fold increase relative to the parental strain, achieving a TyOH titer of 63.22 ± 1.72 mg·L^-1^ (**Table 4**), suggesting that the deregulation of the chorismate mutase branch point further increased flux toward the *TAL* branch, rewiring the synthesis of precursors to the expected direction. Finally, the strain TOH3 was constructed by combining the OE of *ARO10* with the insensitive *ARO4* variant (*ARO4\**). This modification affected TyOH production levels, resulting in a final concentration of 273.41 ± 18.50 mg·L^-1^ (**Table 4**), a 23.65-fold increase over the initial background level. This result is consistent with *ARO10* being among the top two highest-confidence upregulation targets identified by the ecFactory process, and highlights the well-documented role of the feedback-resistant *ARO4* allele in relieving the Tyr-mediated inhibition of DAHP synthase. This result underscores the combined contribution of the deregulated precursor supply and the decarboxylation step catalyzed by *ARO10*, which converts 4-HPP into 4-HPA in the TyOH biosynthetic pathway, as the main factors driving flux towards its accumulation. Since Aro10p also decarboxylates phenylpyruvate, the same interventions are expected to increase flux through the Ehrlich pathway from Phe, consistent with the concomitant accumulation of 2PE in TOH3 (371.31 ± 1.39 mg·L^-1^).

#### 3.3.2. Elimination of competing carbon, redox, and cofactor drains (B2)

The production of bioactive aromatic compounds in yeast is rooted to its secondary metabolism, which harbors some of the most therapeutically and industrially relevant molecules to be overproduced in a eukaryotic chassis, far beyond the classical fermentative by-products tied to the yeast’s carbon metabolism. However, achieving high titers of these compounds is inherently challenging, as it requires a specific, mechanistically driven strategy that goes beyond simple overexpression of biosynthetic genes. As shown in block 1 (**B1**), the modifications implemented to date have effectively increased TyOH accumulation. Regardless, achieving commercially relevant titers requires eliminating competing carbon sinks and rebalancing redox equivalents and cofactor usage, modulations whose targets are rarely intuitive and difficult to predict. To address this, a series of five cumulative modifications was introduced into the TOH3 chassis, each targeting a different metabolic bottleneck. The resulting strains are described sequentially below, with TyOH titers reported at the end of the fermentation (**Table 4**).

Based on the model’s results, we have increased the TyOH titer by approximately 24-fold so far. Given this, we followed its recommendations and implemented *LPP1* downregulation (KD) by replacing its native promoter with a weak constitutive promoter via CRISPR-Cas9-mediated editing, as detailed in the Materials and Methods section, yielding strain TOH4. The KD of *LPP1* increased the TyOH concentration to 331.83 ± 8.25 mg·L^-1^ (**Table 4**), representing a 28.7-fold improvement over the starting laboratory strain, without any measurable negative impact on final biomass (**Table 4**). A consistent result indicating that reduced *LPP1* activity did not compromise cellular fitness but rather redistributed metabolic resources more efficiently toward the target pathway, reducing the carbon drain on phosphorylated shikimate derivatives and driving a net gain of carbon into this pathway. Furthermore, it reduced the energetic burden of lipid biosynthesis, theoretically freeing ATP and redox equivalents that the cell can redirect toward growth and TyOH accumulation.

Next, we removed the *ALD5* and *MAE1* alleles, generating the TOH5 and TOH6 strains, respectively. TyOH production reached 391.05 ± 15.10 mg·L^-1^ and 347.58 ± 10.06, in TOH5 and TOH6, representing increases of 33.8- and 30.1-fold over the WT, respectively (**Table 4**). The KO of *ALD5* resulted in a further increase in TyOH titer relative to both the parental strain and the previous strain in the series, consistent with the expected reduction in diversion of 4-HPAA toward the competing 4-HPA-forming route discussed previously. However, quantifying residual 4-HPA levels before and after this modification would be useful for properly assessing the true impact of *ALD5* knockout on the pathway, thereby allowing a direct comparison of its functional efficiency with that of the well-characterized *feaB* gene in *E. coli*. The absolute titer in TOH6 was, in contrast, lower than that achieved in TOH5, indicating that this modification did not further improve, and in fact partially offset, the gain obtained through *ALD5* KO. Notably, this decrease in titer was accompanied by improved strain growth, with TOH6 reaching a higher final OD_600_ than TOH5. As a result, the OD_600_-normalized specific titer was also lower than that of TOH5, reflecting the combined effects of reduced absolute production and increased biomass accumulation. Nevertheless, this trade-off should not necessarily be regarded as unfavorable. Given the cumulative nature of the strain construction, a modification that improves overall fitness at this intermediate stage, even at the expense of a comparatively smaller reduction in titer, may still be advantageous for the strain’s robustness as further modifications are layered on top.

Following the removal of competing carbon and redox drains, *ZWF1* was downregulated (TOH7 strain) by replacing its native promoter. Subsequently, *ARO1* was OE (TOH8) by replacing its native promoter with the *PDC1* strong promoter, rather than introducing additional copies of the gene at a specific integration site, thereby preserving the gene copy number while substantially increasing its expression level.

Both modifications resulted in a reduction in the final biomass (**Table 4**), followed by OD_600_ measurements, compared to the previous strain in the series, which was, although accompanied by an increase in the normalized yield (mg_TyOH_·g_CDW_^-1^), indicating the enhanced production was not merely proportional to biomass, but reflecting a genuine improvement in per-cell biosynthetic capacity. In the case of *ZWF1*, derepressing its expression resulted in a TyOH titer of 328.83 ± 10.38 mg·L^-1^, whereas OE of *ARO1* reported a TyOH titer of 361.86 ± 27.87 mg·L^-1^ (**Table 4**). This represents a relatively low amount compared with previous titers, suggesting a poor increase or, in fact, a slight stagnation. However, when examining the yields per biomass, we see that the amount of TyOH accumulated is much higher than before, with yields of 397.92 ± 7.77 and 228.24 ± 23.53 mg·g_CDW_^-1^, respectively (**Table 5**). Given this situation, the result in TOH7 is consistent with the expected trade-off discussed in the predictive pipeline, speculating that the reduced flux through the oxidative branch of the PPP limits NADPH regeneration, which can constrain growth while simultaneously favoring carbon-conserving flux toward E4P and, ultimately, TyOH, resembling the growth-production coupling strategy previously exploited through NADPH-limiting *ZWF1* deletion in Tyr-producing strains (Ding et al., 2024; Verho et al., 2003).

**Table 5.** Product yields on substrate and on biomass across the TOH strain series.

| **Strain** | **S (g·L<sup>-1</sup>)** | **Y<sub>EtOH/S</sub><br>(g<sub>EtOH</sub>·g<sub>S</sub><sup>-1</sup>)** | **Y<sub>Gly/S</sub><br>(g<sub>Gly</sub>·g<sub>S</sub><sup>-1</sup>)** | **Y<sub>TyOH/S</sub><br>(mg<sub>TyOH</sub>·g<sub>S</sub><sup>-1</sup>)** | **Y<sub>2PE/S</sub><br>(mg<sub>2PE</sub>·g<sub>S</sub><sup>-1</sup>)** | **Y<sub>TyOH/X</sub><br>(mg<sub>TyOH</sub>·g<sub>X</sub><sup>-1</sup>)** | **Y<sub>2PE/X</sub><br>(mg<sub>2PE</sub>·g<sub>X</sub><sup>-1</sup>)** |
|---|---|---|---|---|---|---|---|
| **WT** | 65.12 ± 0.29<sup>cd</sup> | 0.35 ± 0.01<sup>c</sup> | 0.08 ± 0.00<sup>abcd</sup> | 0.18 ± 0.01<sup>a</sup> | 0.52 ± 0.05<sup>a</sup> | 10.01 ± 0.35<sup>a</sup> | 29.50 ± 2.15<sup>a</sup> |
| **TOH1** | 62.14 ± 3.16<sup>bc</sup> | 0.41 ± 0.04<sup>d</sup> | 0.10 ± 0.01<sup>cd</sup> | 0.54 ± 0.03<sup>a</sup> | 0.71 ± 0.05<sup>a</sup> | 27.18 ± 0.35<sup>a</sup> | 36.25 ± 3.17<sup>a</sup> |
| **TOH2** | 67.75 ± 0.59<sup>cde</sup> | 0.25 ± 0.01<sup>b</sup> | 0.07 ± 0.00<sup>a</sup> | 0.93 ± 0.03<sup>a</sup> | 0.96 ± 0.07<sup>a</sup> | 52.40 ± 1.09<sup>b</sup> | 53.95 ± 4.69<sup>a</sup> |
| **TOH3** | 52.88 ± 1.53<sup>b</sup> | 0.41 ± 0.02<sup>d</sup> | 0.13 ± 0.00<sup>e</sup> | 5.17 ± 0.37<sup>b</sup> | 7.02 ± 0.27<sup>c</sup> | 222.43 ± 1.36<sup>c</sup> | 302.32 ± 6.14<sup>c</sup> |
| **TOH4** | 59.57 ± 2.29<sup>bc</sup> | 0.32 ± 0.00<sup>c</sup> | 0.12 ± 0.01<sup>e</sup> | 5.57 ± 0.08<sup>b</sup> | 6.99 ± 0.43<sup>c</sup> | 266.98 ± 0.61<sup>e</sup> | 334.77 ± 17.32<sup>cd</sup> |
| **TOH5** | 76.28 ± 6.29<sup>efg</sup> | 0.25 ± 0.02<sup>b</sup> | 0.09 ± 0.00<sup>bcd</sup> | 5.14 ± 0.34<sup>b</sup> | 8.71 ± 0.66<sup>d</sup> | 316.55 ± 2.71<sup>f</sup> | 536.22 ± 28.54<sup>f</sup> |
| **TOH6** | 85.98 ± 0.38<sup>g</sup> | 0.27 ± 0.03<sup>b</sup> | 0.08 ± 0.00<sup>abc</sup> | 4.04 ± 0.10<sup>b</sup> | 7.49 ± 0.17<sup>c</sup> | 252.87 ± 1.25<sup>de</sup> | 468.56 ± 20.37<sup>e</sup> |
| **TOH7** | 38.55 ± 5.05<sup>a</sup> | 0.25 ± 0.01<sup>b</sup> | 0.08 ± 0.01<sup>abc</sup> | 8.64 ± 1.29<sup>b</sup> | 6.57 ± 0.36<sup>c</sup> | 397.92 ± 7.77<sup>g</sup> | 305.34 ± 28.90<sup>c</sup> |
| **TOH8** | 39.92 ± 3.17<sup>a</sup> | 0.19 ± 0.02<sup>a</sup> | 0.07 ± 0.01<sup>ab</sup> | 5.39 ± 0.87<sup>c</sup> | 9.05 ± 0.85<sup>c</sup> | 228.24 ± 23.53<sup>cd</sup> | 383.89 ± 9.30<sup>d</sup> |
| **TOH9** | 67.13 ± 4.73<sup>cde</sup> | 0.33 ± 0.01<sup>c</sup> | 0.10 ± 0.02<sup>d</sup> | 8.29 ± 0.32<sup>b</sup> | 5.16 ± 0.30<sup>d</sup> | 506.85 ± 5.17<sup>h</sup> | 316.13 ± 25.38<sup>c</sup> |
| **TOH10** | 80.45 ± 3.54<sup>fg</sup> | 0.19 ± 0.00<sup>a</sup> | 0.10 ± 0.00<sup>cd</sup> | 9.83 ± 0.88<sup>c</sup> | 2.70 ± 0.07<sup>b</sup> | 738.70 ± 9.74<sup>j</sup> | 204.03 ± 18.05<sup>b</sup> |
| **TOH11** | 73.94 ± 2.23<sup>def</sup> | 0.37 ± 0.01<sup>cd</sup> | 0.10 ± 0.00<sup>d</sup> | 12.22 ± 0.19<sup>d</sup> | 3.68 ± 0.22<sup>b</sup> | 650.23 ± 8.81<sup>i</sup> | 196.10 ± 15.61<sup>b</sup> |

Final glucose consumed (S) is expressed in g·L<sup>-1</sup>. Yields of ethanol (EtOH) and glycerol (Gly) are expressed on substrate (g<sub>p</sub>·g<sub>S</sub><sup>-1</sup>), while TyOH and 2PE are expressed on substrate (g<sub>p</sub>·g<sub>S</sub><sup>-1</sup>) and biomass (g<sub>p</sub>·g<sub>X</sub><sup>-1</sup>). Within each column, values sharing the same letter are not significantly different (one-way ANOVA followed by Tukey's post hoc test, *p* < 0.05). Letters apply to each column independently and should not be compared across columns. Values are mean ± SD (n = 3).

In the case of *ARO1* overexpression (TOH8), the observed reduction of final biomass may instead reflect the metabolic burden associated with substantially increasing the expression of *ARO1* at its native locus. Unlike the other OE targets discussed in this work, Aro1p is an unusually large, pentafunctional polypeptide (1588 amino acids, ∼ 174.5 kDa) resulting from the evolutionary fusion of five catalytic domains homologous to five monofunctional *E. coli* enzymes (Duncan et al., 1987), several of which require dedicated cofactors, including NADPH for its shikimate dehydrogenase activity and ATP for its shikimate kinase activity. The synthesis, folding, and cofactor-dependent activity of such a large multidomain enzyme is expected to impose a substantially higher demand on cellular resources (ribosomal capacity, chaperones, ATP, and NADPH) than that of smaller, monofunctional enzymes, potentially limiting the extent to which increased transcription alone can translate into proportionally higher levels of functional enzyme. This metabolic cost may be further intensified by the concurrent NADPH limitation imposed by *ZWF1* downregulation in the preceding strain (TOH7), as both *ARO1* and the upstream PPP compete for the same pool of reduced cofactors (Bertels et al., 2021). Despite this, the performance of the TOH8 strain was about 375.42 ± 30.66 mg·L^-1^ of TyOH (**Table 4**), representing a 1,14-fold increase relative to the TOH7. Thus, we can conclude that the trade-off was generally favorable for the flux towards TyOH.

#### 3.3.3. Boosting the committed flux: maximizing the 4-HPAA-to-TyOH conversion (B3)

Building upon the enhanced precursor availability and diminished competing flux achieved in the preceding strains, this final set of modifications focused on the terminal segment of the Ehrlich pathway: eliminating the last major competing branch at the PPA node (*PHA2*), maximizing the specific conversion capacity of the terminal 4-HPAA conversion (*ADH6, ADH7*), and reinforcing the Tyr-committed step itself through a feedback-insensitive route (*MtPDH1*).

Given that the increased flux of precursors generated in our cell factory also affects the synthesis of another higher alcohol derived from this pathway, namely 2PE, we decided to deregulate the *PHA2* allele to reduce flux towards this competing pathway, in which PPA diverts away from the Tyr branch toward Phe synthesis.

*PHA2* was KD rather than fully deleted by replacing the native promoter with a weak constitutive promoter to reduce flux through this competing branch while minimizing the risk of generating a Phe auxotroph, which is undesirable for process sustainability and scale-up, as amino acid supplementation would increase process cost. This modification (TOH9) resulted in 555.78 ± 20.23 mg·L^-1^ TyOH, a 114.24-fold increase relative to the WT, consistent with the partial redirection of prephenate-derived flux toward the tyrosine-committed branch (**Table 4**). Growth of TOH9 was not affected by the KD. Unlike the decrease observed in the previous engineered versions, growth improved slightly, reaching the same OD_600_ values as the parental strain, suggesting that the residual *PHA2* activity was sufficient to sustain normal Phe biosynthesis under these conditions. To further assess the effect of *PHA2* downregulation on flux redirection away from the competing Phe branch, extracellular 2PE concentration was quantified across the strain series. The KD of *MAE1* in TOH6 did not compromise growth, whereas strains TOH7 and TOH8 showed a reduced final biomass. Consistently, total 2PE accumulation in these two strains was already lower than in TOH6, regardless of *PHA2* status, since 2PE accumulation scales with overall biomass and Ehrlich-pathway flux. This confounds any direct comparison of 2PE levels between TOH9 (*PHA2* KD) and the immediately preceding strains, as a reduction in 2PE level cannot be unambiguously attributed to *PHA2* downregulation rather than to the lower biomass already present in its parental background. For this reason, TOH6 (*MAE1* KD), the last strain in the series that showed unaffected growth, was used instead as the most appropriate reference point to isolate the specific contribution of the reduction in *PHA2* expression, since both TOH6 and TOH9 reached comparable final biomass. Under this comparison, 2PE titer in TOH9 was reduced from 643.56 ± 11.04 mg·L^-1^ to 347.60 ± 35.01 mg·L^-1^ (**Table 3**), which represents approximately half relative to TOH6, providing direct evidence that the KD successfully redirected flux away from the competing branch and, by extension, toward the *TAL* branch.

The terminal step of TyOH biosynthesis, the reduction of 4-HPAA to TyOH, can be catalyzed by several alcohol dehydrogenases using either NADH or NADPH. To reinforce this step, we co-OE *ADH6* and *ADH7* (TOH10 strain), which encode NADPH-dependent enzymes (Larroy, Parés, et al., 2002). Combined OE of both ADH isoforms yielded 788.73 ± 36.84 mg·L^-1^ TyOH, a 68.23-fold increase relative to the WT strain, reaching our highest absolute concentration of TyOH at that time (**Table 4**). Finally, as discussed above, the *M. truncatula* ortholog *MtPDH1,* a cytosolic prephenate dehydrogenase insensitive to Tyr-mediated feedback inhibition, was integrated at a specific genomic locus without altering native *TYR1* expression, thereby generating the final strain of the series (TOH11). This modification increased the final amount of TyOH produced to 902.97 ± 19.43 mg·L^-1^ (**Table 4**).

Altogether, the cumulative modifications raised the TyOH titer of the final strain, TOH11, more than 80-fold relative to the parental strain, reaching the highest concentration obtained in this work (**Figure 2**). This increase was achieved with only a moderate impact on biomass in most strains, as reflected by the biomass values (**Table 4**). Notably, the improvement in TyOH was accompanied by a progressive accumulation of 2PE, indicating that part of the carbon flux was still being diverted towards this competing product.

**Figure 2.**
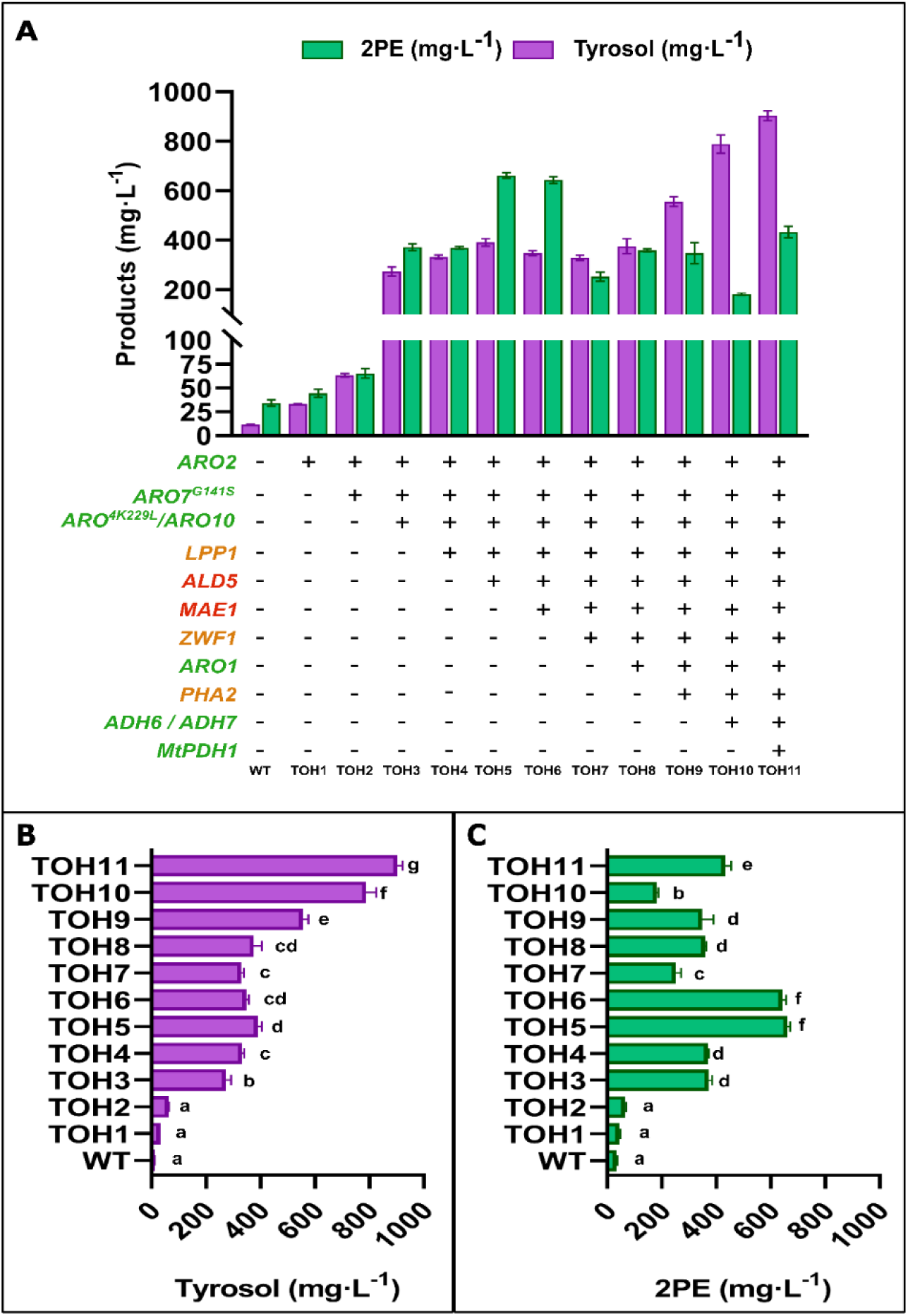
Final titers of TyOH and 2PE achieved by the TOH series compared with the parental strain. Cells were cultured in synthetic defined medium with 80 g·L-^1^ of glucose as the sole carbon source. Cultures were sampled after 7 days of growth to quantify TyOH. **(A)** TyOH and 2PE titers (mg·L^-1^) of the full set of engineered strains, following model-driven predictions and rationally engineered targets. OE targets are shown in green, KD in orange, and finally, KO in red. (**B)** Statistical comparison of TyOH titers among strains. **(C)** Statistical comparison of 2PE titers among strains. In **(B)** and **(C)**, strains sharing a letter do not differ significantly, whereas strains with different letters do. Letters were assigned independently for each panel in ascending order of mean production, so that “a” denotes the group with the lowest production and the last letter (g in **B**, f in **C**) denotes the group with the highest. Significance was assessed using one-way ANOVA followed by Tukey’s HSD post hoc test (p < 0.05). Data are means of n = 3 independent replicates; error bars indicate standard deviation.

### 3.4. Carbon distribution, growth and product yields

The increase in titers does not by itself indicate how efficiently carbon is converted into the targeted product. To address this, we quantified glucose consumption and the formation of ethanol and glycerol across the strain series, and calculated product yields of TyOH and 2PE (**Table 5**). These data were used to (i) examine how the successive modifications redistributed carbon between TyOH, 2PE, and by-products, and (ii) estimate how close TOH11 is to the theoretical limits through a production envelope analysis.

The fermentative profile of the WT strain was compared with the complete pool of engineered strains, since constructing a robust cell factory requires the host’s physiological performance to remain minimally invariant with respect to the targeted product. Glycerol and ethanol yields remained largely unchanged across the series (**Table 5**), indicating that the modifications did not substantially alter by-product formation. Absolute ethanol measurements, even in the WT strain, remained below the expected absolute concentration (43 g·L^-1^) given the initial glucose availability of 86 g·L^-1^.

This implies that a huge portion of carbon is being redirected towards the synthesis of other unmeasured compounds. However, it is worth noting that glucose was not fully depleted by the end of cultivation in most strains, with the notable exception of TOH5 and TOH6, which achieved near-complete glucose consumption (**Table 5**).

In contrast, the yields of TyOH and 2PE varied considerably. Previously, we observed that the absolute concentration of TyOH reached 902.968 ± 19.43 mg·L^-1^, an 80-fold increase over the parental strain (**Table 4**). The first modifications (TOH1–2) raised Y_TyOH/S_ from 0.18 in the WT to 0.93 mg·gS^-1^, while 2PE yields remained below 1 mg·gS^-1^. Nevertheless, the WT produced markedly more 2PE than TyOH from the beginning. A pronounced step change occurred with TOH3, in which Y_TyOH/S_ and Y_2PE/S_ increased 5.6- and 7.3-fold, respectively, suggesting that this modification enhanced carbon flux through a node shared by both branches upstream to the PPA node, before diverging via *PHA2* (toward Phe and 2PE) and *TYR1* (toward Tyr and TyOH) (**Table 5**). Subsequent strains maintained both yields within a similar range (4.0–9.1 mg_TyOH_·gS^-1^), with the specific yield of 2PE generally exceeding TyOH, since the upstream push-pull and flux-redirecting modifications introduced up to this point reinforce the shared trunk pathway feeding both branches without yet specifically redirecting flux through Tyr. This trait was truncated only upon the introduction of the *PHA2* knockdown (TOH9), which substantially reversed this balance. Notably, this occurred without greatly impacting growth fitness. From TOH9 onward, Y_2PE/S_ fell to 2.70-3.68 mg·gS^-1^, whereas TyOH yield continued to increase steadily, reaching a final specific yield of 12.22 ± 0.19 mg⁻¹·gS⁻¹, so that the 2PE:TyOH ratio decreased from 2.9 in the parental strain to 0.30 in the final strain. Overall, Y_TyOH/S_ increased approximately 68-fold relative to the WT strain. Yields per unit of biomass followed the same trend, with Y_TyOH/X_ increasing from 10.01 ± 0.35 mg·gS⁻¹ in WT to 650.23 ± 8.81 mg·gS⁻¹ in TOH11 (**Table 5**), although the maximum value peaked in TOH10 (738.70 ± 9.74 mg·gS⁻¹). These findings were obtained in a laboratory background with an inherent bias towards the Phe branch. Industrial *S. cerevisiae* strains, including those used in winemaking, show considerable strain-dependent variability in the production of Tyr-derived metabolites (Cordente et al., 2021; Guerrini et al., 2018), suggesting that some of them could offer a more suitable intrinsic flux distribution towards TyOH and a better starting point for the same engineered purpose.

#### 3.4.1. Growth–production trade-off: FBA-derived production envelopes

Successive engineering steps reshaped the predicted trade-off between growth and TyOH production while initially leaving the theoretical yield ceiling unchanged. To examine this progression, we simulated a production envelope for each mutant strain and compared it with the corresponding experimental measurement (**Figure 3**). Each envelope’s upper boundary defines the maximum theoretical TyOH yield at a given growth rate. The parental strain (WT) displayed the largest feasible area, providing a reference for following how successive modifications constrained biomass formation and TyOH production.

**Figure 3.**
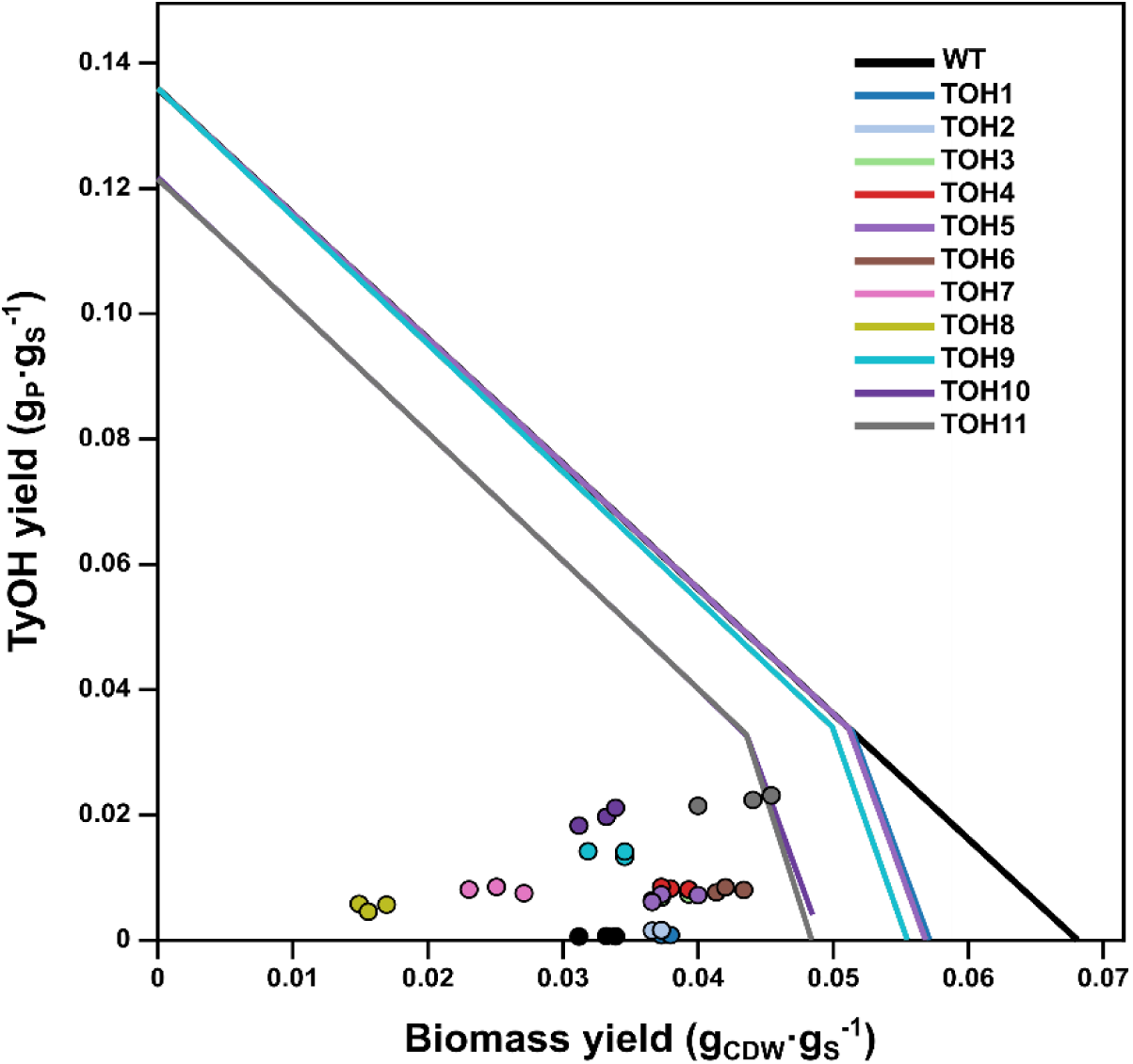
Production envelopes and experimental yields for the TyOH-overproducing *S. cerevisiae* strains. Lines show the predicted TyOH-growth trade-off for each tested strain; dots show experimental yields in flask cultures. The WT and mutants up to TOH9 exhibit similar behavior, with a maximum theoretical yield of approximately 0.136 g·g^-1^, while their maximum biomass yields progressively decline. TOH11 and TOH10 have a lower TyOH ceiling of approximately 0.121 g·g^-1^, indicating a constraint at the final reduction step. Although all experimental yields fall below the predicted barrier, tyrosol production increases across the strain series, and TOH9, TOH10, and TOH11 lie proportionally closest to their respective limits.

Across the first nine strains, from WT through TOH7, the maximum TyOH yield remained close to the theoretical ceiling of 0.136 g·gS^-1^. The modifications therefore altered the feasible combinations of growth and production while preserving the maximum attainable yield. The early overexpression targets link two connected pathways: the shikimate pathway (*ARO2*, *ARO7*, *ARO4*, *ARO1*), which supplies chorismate, the shared precursor for all aromatic amino acids, including Tyr, and the Ehrlich pathway (*ARO10*, *ADH7*), which converts Tyr directly into TyOH. Combined with the KD and KO steps, these edits progressively narrowed the feasible space towards higher TyOH-to-biomass ratios without increasing the stoichiometric yield ceiling (**Figure 3**).

Overexpression of *ARO1* is particularly important in this context, as CHA is the shared precursor for Phe, Trp, and Tyr biosynthesis, for growth via protein synthesis, and for TyOH, which draws on the same pool. By increasing CHA supply, *ARO1* OE prevents this shared precursor step from becoming rate-limiting as subsequent edits divert an increasing share of Tyr flux toward TyOH. Maintaining *ARO1* flux, therefore, keeps this bottleneck open as the other edits accumulate. A modest decrease in the maximum tyrosol yield, to approximately 0.122 g·g^-1^, first occurs in TOH10 and persists in TOH11, coinciding with the overexpression of *ADH6* and *ADH7*. Across the strain series, maximum biomass yield declines gradually from 0.068 gX·gS^-1^ in the WT to 0.048 gX·gS^-1^ in TOH11, reflecting the cumulative metabolic burden imposed by the KD/KO modifications.

Comparison with the experimental flask data reveals a clear shift in how closely realized performance tracks the predicted envelope. In the early strains, from the WT through TOH7, measured TyOH yields remain below the predicted trade-off curve, indicating that only a small fraction of the theoretical growth-production trade-off is accessed under batch conditions. From TOH8 onward, although the predicted maximum is lower, the measured yields lie proportionally closer to this predicted limit. The highest-producing laboratory strains, TOH9, TOH10, and TOH11, therefore, achieve a substantial fraction of their reduced theoretical maximum. Thus, this pattern suggests that the cumulative edits improved TyOH titer not only in absolute terms but also by bringing the strain’s actual yield closer to the model-predicted potential for each genotype.

Taken together, these results validate the overall push-pull and flux-redirection strategy pursued in this work and establish a solid genetic background upon which to carry out further process-level optimization, including scale-up to bioreactor cultivation and detailed characterization of growth kinetics (specific growth rate, biomass, and substrate yields or productivity, among others) under more tightly controlled conditions, establishing our cell factory as one of the top-ranked productive strains in small-scale conditions.

#### 3.4.2. Growth kinetics of the engineered strains

To formally quantify the growth burden associated with each modification, all strains were cultured in 48-well microplates under the same conditions as those used for the shake flasks. The growth curves were fitted to the Gompertz model (**Table 6**) to determine the kinetic parameters for the complete strain series. All strains displayed the typical sigmoid yeast growth curve and closely followed the growth pattern observed in the WT strain (**Figure S1A**), with no significant differences in μ_max_ or apparent maximum OD_600_ across the series. The main burden was an extended lag phase, which increased from ∼4 h in the WT to ∼20 h in the final strain TOH11, reaching 23–28 h in TOH8 and TOH9 (**Figure S1B**). However, this burden was not cumulative, as the definitive strain showed a maximum growth similar to that of the WT. Accordingly, the AUC decreased most in TOH8 and TOH9 (to 72 % and 64 % of the WT value, respectively) and moderately in TOH11 (85 %), but no strain lost more than ∼ 37 % of the WT value. Shake-flask cultures (**Table 5**) were broadly consistent with these trends up to TOH6, but from TOH7 onward, growth was poorer than the microplates suggested, and some strains behaved differently between formats (**Figure S1A**), possibly reflecting differences in aeration and oxygen transfer. Overall, the modifications primarily delayed the onset of growth rather than reducing the exponential growth rate, which may reflect an adaptation cost rather than a permanent loss of growth capacity. The factors underlying this delay (for example, nutrient availability, pH, agitation, or the osmotic stress imposed by the high glucose concentration) remain to be investigated.

**Table 6.**
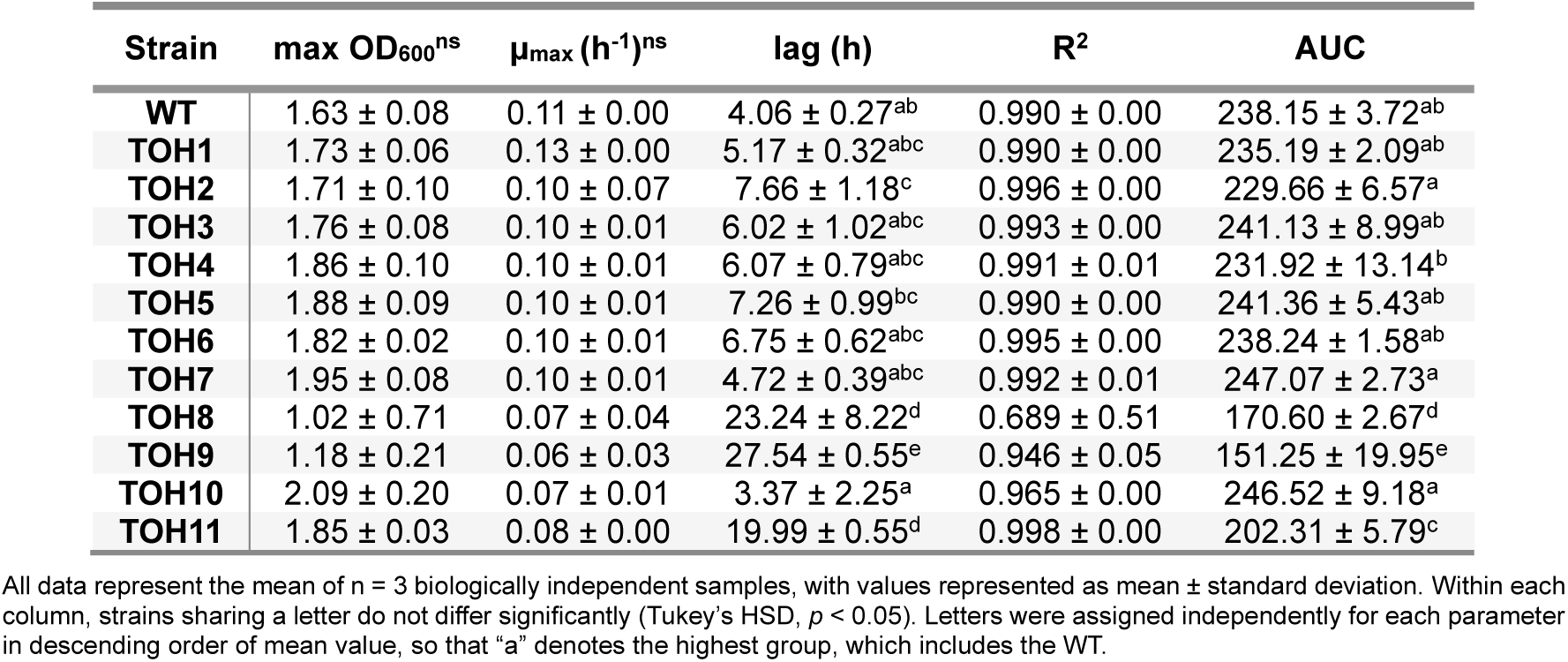
Comparison of growth kinetic parameters across the engineered strains after 96 h. . For each strain, the apparent maximum **OD_600_**, the **μ_max_**, the lag time, the goodness of fit (**R²**), and the area under the curve (**AUC**) at the end point are reported.

### 3.5. Batch-scale validation and dynamic flux balance analysis of the final TOH strain

While the shake flask and microplate-based characterization presented above in sections allowed the relative performance of the complete strain series to be compared under controlled and reproducible conditions, these small-scale formats present inherent limitations for a definitive assessment of production performance, including limited aeration, pH and temperature variations, restricted sampling frequency, and the absence of direct process monitoring. To address these limitations and validate the production phenotype of the final engineered strain under more representative conditions, the definitive TOH strain (TOH11) and the parental strain were characterized in parallel in batch bioreactors (Applikon) in triplicate at a larger working volume (0.25 L). The more frequent and precise sampling enabled a better determination of the growth profile, including the maximum specific growth rate of each strain, calculated by linear regression of ln(OD_600_) versus time over the exponential phase (**Figure S2**). This allowed time-resolved quantification of biomass accumulation (OD_600_ and CDW), glucose consumption, and TyOH production.

Beyond providing a direct, scaled-up comparison between the parental and TOH11 strains, this batch dataset also served as the basis for parameterizing a dynamic flux balance analysis (dFBA) model. Comparing these dynamic model predictions with experimentally observed batch kinetics provides additional validation of the metabolic model underlying the ecFactory-derived target identification presented earlier, beyond the single-timepoint or steady-state comparisons performed so far.

#### 3.5.1. Batch-scale TyOH overproduction

As a first approximation of what a larger production volume would entail, 0.25 L batch cultures in SD80 medium were performed for both the parental and the TOH11 strain. This experiment aimed, on the one hand, to confirm whether TyOH production would decrease or otherwise be affected by the increased surface area available relative to shake flask cultivation, and, on the other hand, to determine whether growth and glucose consumption dynamics would be similar to those observed previously **(Figure 4)**.

**Figure 4.**
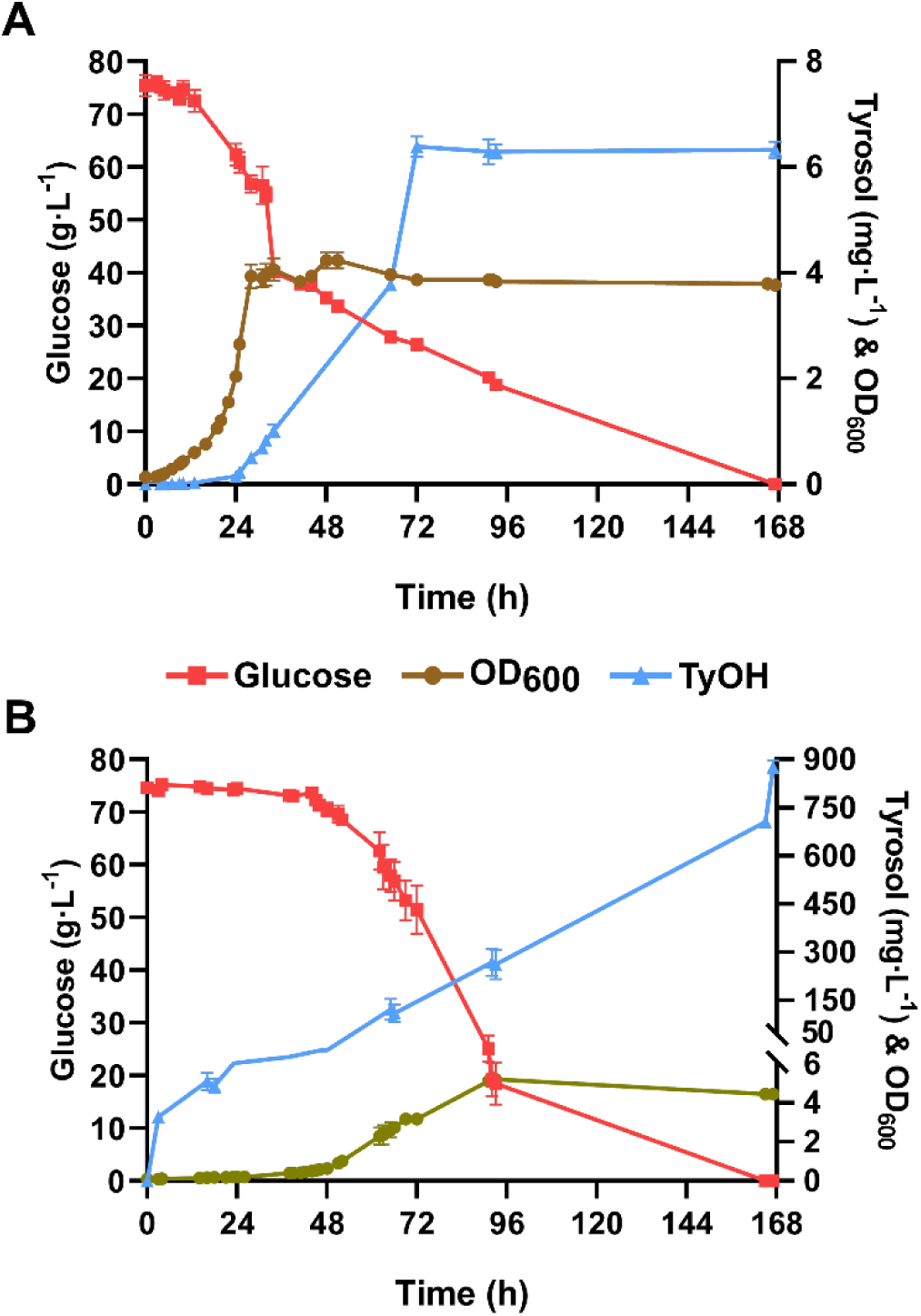
TyOH production assessment in batch-scale bioreactors. A production comparison is made between the WT and the TOH11 strain. Glucose and TyOH are expressed in g·L^-1^, while biomass was recorded as OD_600_. **A.** Parental strain growth and product formation over time. **B.** TOH11 cell factory growth profile and product formation over time. All data represent the mean of n = 3 biologically independent samples, with values represented as mean ± standard deviation.

The WT strain displayed faster growth and a higher initial glucose consumption rate. Both cultures were maintained for a total of 7 days, with approximately 28 g·L^-1^ of residual substrate remaining in the parental strain at 72 hours, compared with 51 g·L^-1^ for TOH11 at the same cultivation point.

Under these conditions, the parental strain produced approximately 6.38 ± 0.19 mg·L^-1^ of TYOH, which was very similar to the value quantified in the shake-flask cultivations (**Figure 4A**). In comparison, the engineered strain (TOH11) required 24 h longer to consume the same amount of glucose as the WT, reaching a comparable level of consumption only by around 96 h, compared to the WT’s 72 h (**Figure 4B**). However, given that our cell factory had shown during previous production runs that TyOH accumulation increased over time, the batch culture was deliberately left to run longer to maximize TyOH production.

Despite this slower growth, TyOH production in the final strain was 876.91 ± 20.37 mg·L^-1^, approximately 140-fold higher than in the WT (**Figure 4B**). Notably, TyOH accumulation continued to increase progressively throughout cultivation, without being tightly coupled to either glucose consumption or biomass accumulation, suggesting that part of the TyOH is produced non-growth-associated, increasing the final amounts during the stationary phase, once active cell division has largely ceased, and also suggesting that TyOH is released into the supernatant once the stationary phase is reached.

Overall, these results confirm that the production phenotype of the final strain was preserved upon scaling up from shake flask to batch cultivation at a larger working volume, with a relative improvement in TyOH yield, which led to more than 100-fold higher TyOH production under batch conditions, regardless of vessel size or format (**Table 7**). At this larger scale, the compromise between growth and TyOH production was even more favorable than in shake flasks, with higher titers achieved at comparable biomass levels.

**Table 7.** Batch process parameters for TyOH production in the WT and the TOH11 cell factory.

|  | 72 h |  | 167 h |  |
|---|---|---|---|---|
| **Parameters** | **WT** | **TOH11** | **WT** | **TOH11** |
| **OD<sub>600</sub>** | 3.83 ± 0.12 | 3.18 ± 0.29 | 3.77 ± 0.06 | 4.47 ± 0.11 |
| **CDW (g·L<sup>-1</sup>)** | 0.99 ± 0.03 | 0.73 ± 0.06 | 0.90 ± 0.03 | 1.01 ± 0.025 |
| **TyOH (mg· L<sup>-1</sup>)** | 6.39 ± 0.19 | 121.11 ± 32.84 | 6.33 ± 0.16 | 876.91 ± 20.37 |
| **Yield (mg<sub>TyOH</sub>·g<sub>CDW</sub><sup>-1</sup>)** | 7.13 ± 0.38 | 148.89 ± 25.95 | 7.06 ± 0.15 | 869.22 ± 33.58 |
| **Yield (mg<sub>TyOH</sub>·g<sub>S</sub><sup>-1</sup>)** | 0.08 ± 0.00 | 5.52 ± 0.91 | 0.08 ± 0.00 | 11.44 ± 0.27 |

Parameters were calculated at 72 h and at the end of batch cultivation (167 h). Biomass was monitored by OD<sub>600,</sub> and cell dry weight (CDW) was calculated for each strain. Net TyOH titer (mg·L<sup>-1</sup>), product yield relative to biomass (Y<sub>P/CDW</sub>), and relative to consumed substrate (Y<sub>P/S</sub>) are reported.

Product yields relative to biomass obtained for TOH11 with respect to the WT showed that TyOH increased 23-fold (Y_P/CDW_) at 72 h, whereas the yield relative to consumed substrate (Y_P/S_) increased 35.1-fold. However, when TOH11 was left in the batch for longer, the total amount of TyOH grew exponentially. At 167 h, the yield relative to biomass reached 134-fold (Y_P/X_), while the yield relative to substrate increased 72.9-fold after glucose was completely depleted.

The combined strategy, including modeling and rational principles, allowed us to develop a strong evidence-based cell factory. Our results were highly positive, and placing them in the context of the existing literature reveals that either our batch and shake-flask yields are competitive with, and in some cases exceed, those reported for other engineered yeast strains (**Table 7)**.

Among engineered *S. cerevisiae* strains, our TOH11 chassis achieved the highest biomass-normalized productivity reported to date (869.22 mg_TyOH_·gX^-1^), surpassing previously reported yeast strains, including the HLF-Dα strain (170.85 mg_TyOH_·gX^-1^) engineered by Guo et al. (2020) and the CEN.PK-based strain of Sun et al. (2025) that achieved 70.59 mg_TyOH_·gX^-1^, or the BY4741 version of Guo et al. (2019) that raised a yield of 126.74 mg_TyOH_·gX^-1,^ despite operating at a lower biomass concentration (**Table 8**). In *E. coli*, higher productivities have been reported: Yang et al. (2018) obtained 719.29 mg_TyOH_·gX^-1^, and Li et al. (2026) got 1030.28 mg_TyOH_·gX^-1^, reflecting the intrinsically higher flux through the aromatic amino acid pathway in that host under the tested conditions (**Table 8**). Nevertheless, the competitive performance of our *S. cerevisiae* chassis at this discontinuous level, together with the inherent advantages of yeast as an industrial host, provides a solid basis for scaling up TyOH production through bioprocess strategies. Notably, the titer-to-biomass yields achieved by our engineered strain in SD80 medium without precursor supplementation and under a comparatively low biomass yield (Y_X/S_) constitute a stringent, industrially relevant benchmark for future optimization efforts.

**Table 8.** Summary of recent reports regarding the production of TyOH from glucose in yeast and bacteria as hosts.

| Host | Strain | Medium | System | X<br>(g·L<sup>-1</sup>) | TyOH<br>(mg·L<sup>-1</sup>) | Y<sub>TyOH/X</sub><br>(mg·g<sub>X</sub><sup>-1</sup>) | Reference |
|---|---|---|---|---|---|---|---|
| *S. cerevisiae* | BY4741 | SC-ura | 0.05 L SF | 1.00 | 126.74 | 126.74 | (Guo et al., 2019) |
|  | HLF-Dα | YPD | 0.05 L SF | 5.43 | 927.68 | 170.85 | (Guo et al., 2020) |
|  | CEN.PK2-1C | YPD | 0.02 L SF | 8.1 | 571.80 | 70.59 | (Sun et al., 2025) |
|  | **BY4743 (TOH11)** | **SD** | **0.03 L SF** | **1.39** | **902.97** | **650.23** | **This study** |
|  |  | **SD** | **0.25 L batch** | **1.01** | **876.91** | **869.22** | **This study** |
| *E. coli* | MG1655 | M9Y | 0.05 L SF | 1.83 | 1316.30 | 719.29 | (Yang et al., 2018) |
|  | MG1655 | LB | 0.05 L SF | 1.72 | 1768.50 | 1030.28 | (Li et al., 2026) |

SF, shake-flask culture; batch, batch culture. Biomass (X) is reflected as CDW (g·L<sup>-1</sup>), volumetric TyOH titer, and TyOH concentration normalized to biomass (Y<sub>TyOH/X</sub>).

#### 3.5.2. Dynamic FBA validation of the ecFactory-predicted TOH11 phenotype

The resulting strain, TOH11, was used to simulate batch cultures under TyOH-producing conditions. Compound accumulation was simulated using dynamic flux balance analysis (dFBA), which extends standard FBA to a time-resolved framework by dividing the cultivation period into short intervals and optimizing at each step (Mahadevan et al., 2002). The model reproduced the overall patterns of substrate consumption, biomass accumulation, and ethanol production during the first 90 h, encompassing exponential growth and the transition towards stationary phase in the final hours. Without fitting the individual curves, the simulation captured the gradual consumption of approximately 56 g·L⁻¹ of substrate, the accumulation of approximately 1.1 g·L^-1^ of biomass, expressed as CDW, and the pronounced ethanol overflow characteristic of fermentative metabolism in batch cultures of *S. cerevisiae*, reaching approximately 24 g·L^-1^ (**Figure 5**).

**Figure 5.**
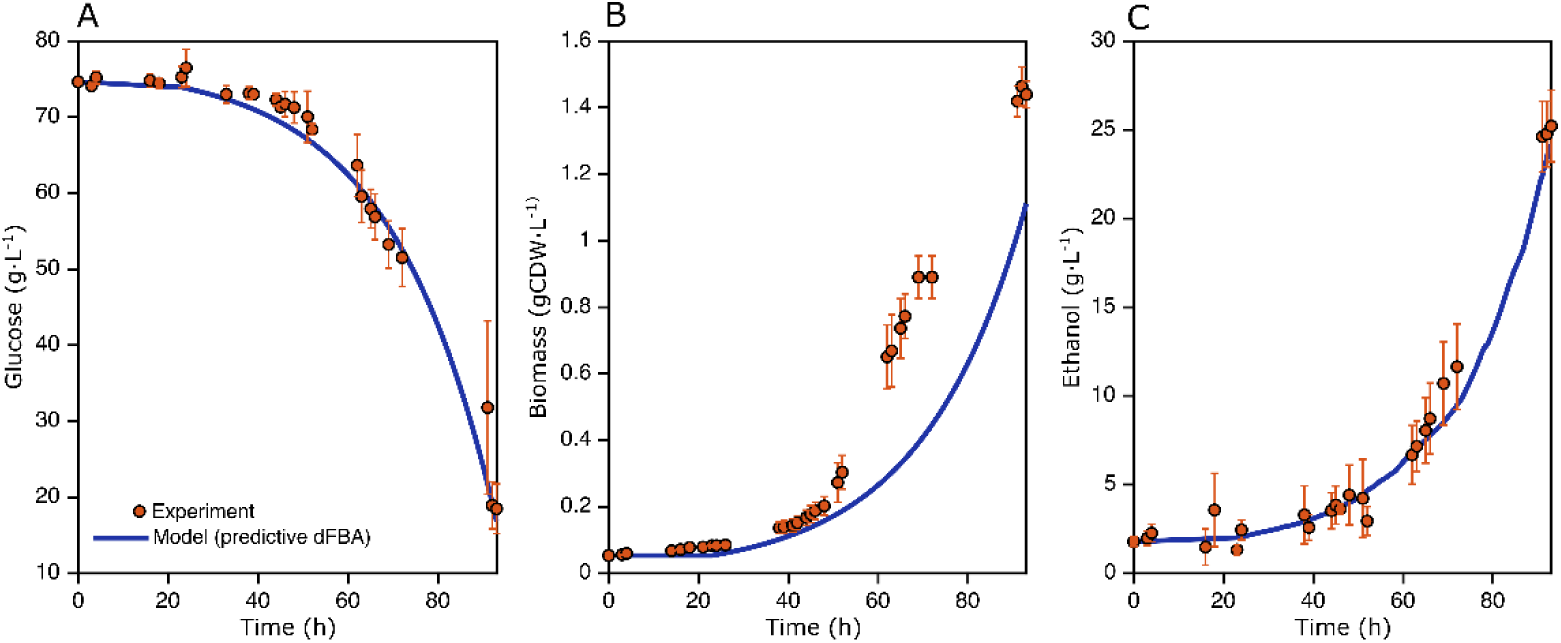
Predictive dynamic FBA of the TOH11 batch culture relative to the synthetic productions assumed by the model. Each panel compares model predictions with triplicate experimental measurements over 93 h. Glucose consumption (**A**), biomass formation (**B**), and ethanol production (**C**) are quantified without fitting, including limited biomass accumulation and pronounced ethanol overflow.

For TyOH formation, the model followed the experimentally observed increasing trend but overestimated the titer before stationary phase by approximately 26-fold, predicting 6.78 g·L^-1^ compared with the measured 0.261 g·L^-1^ (**Figure S3**). This discrepancy is not entirely unexpected, as optimization during dFBA maximizes TyOH production while permitting an *in-silico* protein allocation that cannot be achieved in the current yeast cellular background. In our simulations, protein usage reached approximately 39 % for *ADH7* and 25 % for *ARO2*, illustrating the overestimation of protein allocation to this pathway. Such allocations are biologically implausible, and proteomic data for *S. cerevisiae* indicate that even the most highly expressed glycolytic enzymes under fermentative conditions rarely exceed 3–8 % of total cellular protein (Milo, 2013; Sánchez et al., 2017). The overprediction therefore reflects the absence of enzyme-specific abundance constraints for the introduced pathway enzymes, rather than an error in carbon balance or genotype implementation. Future analyses could incorporate proteomic data to more precisely constrain protein allocation under these conditions and potentially reduce inherent artifacts in ecGEM simulations.

A further biologically distinct source of overprediction concerns the timing of TyOH production rather than its amount. As an aromatic compound derived from the shikimate pathway, TyOH accumulation in flask cultures appears to be concentrated in the stationary phase, once active growth has ceased, consistent with reports that *S. cerevisiae* channels AAA flux toward aromatic alcohol production, including TyOH, under nitrogen-restrictive and growth-arrested conditions (Avbelj et al., 2015). Given that dFBA couples product formation to active biomass accumulation, it is structurally unable to capture this growth-decoupled behavior, thereby reinforcing the effect of enzyme allocation mentioned above and contributing to the overall overestimation of TyOH.

## 4. Conclusions

In this work, a laboratory *Saccharomyces cerevisiae* strain was engineered to overproduce TyOH from a simple, defined medium, combining 13 targets: nine prioritized by an enzyme-constrained genome-scale model (ecFactory) and four selected on rational grounds, all implemented by marker-free CRISPR-Cas9 editing. The model-derived targets, chiefly the combinatorial overexpression of *ARO* genes together with modifications that reduce energy and redox costs, increased the TyOH titer 28.7-fold relative to the parental strain, supporting the predictive value of this approach for identifying push–pull targets. The rationally selected modifications provided complementary gains by favoring the final conversion of 4-HPAA to TyOH, eliminating competing routes (4-HPAA oxidation and 2PE biosynthesis) and redirecting flux through the pentose phosphate pathway towards E4P supply. The final strain reached biomass-specific yields of 650.23 ± 8.81 mg_TyOH_·gX^-1^ in shake flasks and 869.22 ± 33.58 mg_TyOH_·gX^-1^ in 0.5 L batch bioreactors, with maximum titers of 876–901 mg·L^-1^, without extensive optimization of the production process.

Overall, the engineered strain achieved a TyOH titer more than 100-fold higher than that of the WT under equivalent conditions, providing a rational metabolic engineering framework for TyOH overproduction in yeast and laying the foundation for developing a robust cell factory. Since TyOH accumulated mainly in the stationary phase, future work should optimize cultivation conditions to decouple production from biomass formation, for instance, through extended or stationary-phase-oriented cultivation strategies.

## 5. Supporting Information

The supporting material for this study is available free of charge at http://hdl.handle.net/10261/450456. Plasmids and primers used in this work (Tables S1 – S2); ecFactory L3 candidates list containing the selected genes (Table S3); comparison of growth and kinetics for the whole TOH series strains (Figure S1); experimental μ_max_ calculations for WT and TOH11 in batch cultures (Figure S2); Predictive dFBA of the TOH11 batch culture relative to the ecFactory model (Figure S3); raw data and calculation spreadsheets used for data processing and analysis (Excel files, .xls).

## 6. Author Contributions

A.G. contributed to conceptualization, methodology, investigation, formal analysis, data curation, and writing the original draft. F.M. contributed to methodology, investigation (computational modeling), and writing. S.S. contributed to investigation (experimental work). R.B. contributed to supervision. J.M. contributed to funding acquisition and supervision.

## 8. Acknowledgments

This work was funded by MCIN/AEI/10.13039/501100011033, grant numbers PID2022-142748OB-I00 and PID2025-174640OB-I00. AG thank its PRE2020-093597 grant funded by MCIN/AEI. Authors acknowledge the award of the Spanish government MCIN/AEI to the IATA-CSIC as Center of Excellence (CEX2021-001189-S/MICIU/AEI / 10.13039/501100011033).

## References

Adeboye, P. T., Bettiga, M., & Olsson, L. (2017). ALD5, PAD1, ATF1 and ATF2 facilitate the catabolism of coniferyl aldehyde, ferulic acid and p-coumaric acid in Saccharomyces cerevisiae. Scientific Reports, 7(July 2016), 1–13. 10.1038/srep42635

Avbelj, M., Zupan, J., Kranjc, L., & Raspor, P. (2015). Quorum-Sensing Kinetics in Saccharomyces cerevisiae: A Symphony of ARO Genes and Aromatic Alcohols. Journal of Agricultural and Food Chemistry, 63(38), 8544–8550. 10.1021/acs.jafc.5b03400

Becker, J., & Boles, E. (2003). A modified Saccharomyces cerevisiae strain that consumes L-arabinose and produces ethanol. Applied and Environmental Microbiology, 69(7), 4144–4150. 10.1128/AEM.69.7.4144-4150.2003

Bertels, L. K., Murillo, L. F., & Heinisch, J. J. (2021). The pentose phosphate pathway in yeasts–more than a poor cousin of glycolysis. Biomolecules, 11(5). 10.3390/biom11050725

Bisquert, R., Planells-Cárcel, A., Valera-García, E., Guillamón, J. M., & Muñiz-Calvo, S. (2022). Metabolic engineering of Saccharomyces cerevisiae for hydroxytyrosol overproduction directly from glucose. Microbial Biotechnology, 15(5), 1499–1510. 10.1111/1751-7915.13957

Boles, E., De Jong-Gubbels, P., & Pronk, J. T. (1998). Identification and characterization of MAE1, the Saccharomyces cerevisiae structural gene encoding mitochondrial malic enzyme. Journal of Bacteriology, 180(11), 2875–2882. 10.1128/jb.180.11.2875-2882.1998

Borodina, I., & Nielsen, J. (2014). Advances in metabolic engineering of yeast Saccharomyces cerevisiae for production of chemicals. Biotechnology Journal, 9(5), 609–620. 10.1002/biot.201300445

Braus, G. H. (1991). Aromatic amino acid biosynthesis in the yeast Saccharomyces cerevisiae: A model system for the regulation of a eukaryotic biosynthetic pathway. Microbiological Reviews, 55(3), 349–370. 10.1128/mr.55.3.349-370.1991

Britton, J., Davis, R., & O’Connor, K. E. (2019). Chemical, physical and biotechnological approaches to the production of the potent antioxidant hydroxytyrosol. Applied Microbiology and Biotechnology, 103(15), 5957–5974. 10.1007/s00253-019-09914-9

Cadière, A., Ortiz-Julien, A., Camarasa, C., & Dequin, S. (2011). Evolutionary engineered Saccharomyces cerevisiae wine yeast strains with increased in vivo flux through the pentose phosphate pathway. Metabolic Engineering, 13(3), 263–271. 10.1016/j.ymben.2011.01.008

Chen, Y., Gustafsson, J., Tafur Rangel, A., Anton, M., Domenzain, I., Kittikunapong, C., Li, F., Yuan, L., Nielsen, J., & Kerkhoven, E. J. (2024a). Reconstruction, simulation and analysis of enzyme-constrained metabolic models using GECKO Toolbox 3.0. Nature Protocols, 19(3), 629–667. 10.1038/s41596-023-00931-7

Chen, Y., Gustafsson, J., Tafur Rangel, A., Anton, M., Domenzain, I., Kittikunapong, C., Li, F., Yuan, L., Nielsen, J., & Kerkhoven, E. J. (2024b). Reconstruction, simulation and analysis of enzyme-constrained metabolic models using GECKO Toolbox 3.0. Nature Protocols, 19(3), 629–667. 10.1038/s41596-023-00931-7

Cordente, A. G., Nandorfy, D. E., Solomon, M., Schulkin, A., Kolouchova, R., Francis, I. L., & Schmidt, S. A. (2021). Aromatic higher alcohols in wine: Implication on aroma and palate attributes during chardonnay aging. Molecules, 26(16). 10.3390/molecules26164979

Cordente, A. G., Schmidt, S., Beltran, G., Torija, M. J., & Curtin, C. D. (2019). Harnessing yeast metabolism of aromatic amino acids for fermented beverage bioflavouring and bioproduction. Applied Microbiology and Biotechnology, 103(11), 4325–4336. 10.1007/s00253-019-09840-w

Cunha, J. T., Soares, P. O., Baptista, S. L., Costa, C. E., & Domingues, L. (2020). Engineered Saccharomyces cerevisiae for lignocellulosic valorization: a review and perspectives on bioethanol production. Bioengineered, 11(1), 883–903. 10.1080/21655979.2020.1801178

Daniele N. Di, Noce, A., Vidiri, M. F., Moriconi, E., Marrone, G., Annicchiarico-Petruzzelli, M., D’urso, G., Tesauro, M., Rovella, V., & De Lorenzo, A. (2017). Impact of Mediterranean diet on metabolic syndrome, cancer and longevity. Oncotarget, 8(5), 8947–8979. 10.18632/oncotarget.13553

Datta, S., Annapure, U. S., & Timson, D. J. (2017). Different specificities of two aldehyde dehydrogenases from Saccharomyces cerevisiae var. boulardii. Bioscience Reports, 37(2), 1–14. 10.1042/BSR20160529

Deaner, M., & Alper, H. S. (2017). Systematic testing of enzyme perturbation sensitivities via graded dCas9 modulation in Saccharomyces cerevisiae. Metabolic Engineering, 40(February), 14–22. 10.1016/j.ymben.2017.01.012

Ding, N., Yuan, Z., Sun, L., & Yin, L. (2024). Dynamic and Static Regulation of Nicotinamide Adenine Dinucleotide Phosphate: Strategies, Challenges, and Future Directions in Metabolic Engineering. Molecules, 29(15), 3687. 10.3390/molecules29153687

Domenzain, I., Lu, Y., Wang, H., Shi, J., Lu, H., & Nielsen, J. (2025). Computational biology predicts metabolic engineering targets for increased production of 103 valuable chemicals in yeast. Proceedings of the National Academy of Sciences, 122(9). 10.1073/pnas.2417322122

Domenzain, I., Sánchez, B., Anton, M., Kerkhoven, E. J., Millán-Oropeza, A., Henry, C., Siewers, V., Morrissey, J. P., Sonnenschein, N., & Nielsen, J. (2022). Reconstruction of a catalogue of genome-scale metabolic models with enzymatic constraints using GECKO 2.0. Nature Communications, 13(1), 1–13. 10.1038/s41467-022-31421-1

Duncan, K., Edwards, R. M., & Coggins, J. R. (1987). The pentafunctional arom enzyme of Saccharomyces cerevisiae is a mosaic of monofunctional domains. The Biochemical Journal, 246(2), 375–386. 10.1042/bj2460375

Faulkner, A., Chen, X., Rush, J., Horazdovsky, B., Waechter, C. J., Carman, G. M., & Sternweis, P. C. (1999). The LPP1 and DPP1 gene products account for most of the isoprenoid phosphate phosphatase activities in Saccharomyces cerevisiae. Journal of Biological Chemistry, 274(21), 14831–14837. 10.1074/jbc.274.21.14831

Furneisen, J. M., & Carman, G. M. (2000). Enzymological properties of the LPP1-encoded lipid phosphatase from Saccharomyces cerevisiae. Biochimica et Biophysica Acta - Molecular and Cell Biology of Lipids, 1484(1), 71–82. 10.1016/S1388-1981(99)00202-4

Gietz, R. D., & Woods, R. A. (2002). Transformation of yeast by lithium acetate/single-stranded carrier DNA/polyethylene glycol method. Methods in Enzymology, 350(2001), 87–96. 10.1016/S0076-6879(02)50957-5

Gold, N. D., Gowen, C. M., Lussier, F. X., Cautha, S. C., Mahadevan, R., & Martin, V. J. J. (2015). Metabolic engineering of a tyrosine-overproducing yeast platform using targeted metabolomics. Microbial Cell Factories, 14(1), 1–16. 10.1186/s12934-015-0252-2

Gottardi, M., Reifenrath, M., Boles, E., & Tripp, J. (2017). Pathway engineering for the production of heterologous aromatic chemicals and their derivatives in Saccharomyces cerevisiae: bioconversion from glucose. FEMS Yeast Research, 17(4), 1–11. 10.1093/femsyr/fox035

Guerrini, S., Mangani, S., Romboli, Y., Luti, S., Pazzagli, L., & Granchi, L. (2018). Impact of Saccharomyces cerevisiae strains on health-promoting compounds in wine. Fermentation, 4(2), 1–14. 10.3390/fermentation4020026

Guo, W., Huang, Q., Feng, Y., Tan, T., Niu, S., Hou, S., Chen, Z., Du, Z. Q., Shen, Y., & Fang, X. (2020). Rewiring central carbon metabolism for tyrosol and salidroside production in Saccharomyces cerevisiae. Biotechnology and Bioengineering, 117(8), 2410–2419. 10.1002/bit.27370

Guo, W., Huang, Q., Liu, H., Hou, S., Niu, S., Jiang, Y., Bao, X., Shen, Y., & Fang, X. (2019). Rational Engineering of Chorismate-Related Pathways in Saccharomyces cerevisiae for Improving Tyrosol Production. Frontiers in Bioengineering and Biotechnology, 7(JUL), 1–7. 10.3389/fbioe.2019.00152

Guo, X., Wu, X., Ma, H., Liu, H., & Luo, Y. (2023). Yeast: A platform for the production of L-tyrosine derivatives. Yeast, 40(5–6), 214–230. 10.1002/yea.3850

Hazelwood, L. A., Daran, J. M., Van Maris, A. J. A., Pronk, J. T., & Dickinson, J. R. (2008). The Ehrlich pathway for fusel alcohol production: A century of research on Saccharomyces cerevisiae metabolism. Applied and Environmental Microbiology, 74(8), 2259–2266. 10.1128/AEM.02625-07

He, X., & Lu, H. (2025). Evolution and applications of genome-scale metabolic models in yeast systems biology studies. FEMS Yeast Research, 25(August). 10.1093/femsyr/foaf045

Henriques, D., Minebois, R., Mendoza, S. N., Macías, L. G., Pérez-Torrado, R., Barrio, E., Teusink, B., Querol, A., & Balsa-Canto, E. (2021). A Multiphase Multiobjective Dynamic Genome-Scale Model Shows Different Redox Balancing among Yeast Species of the Saccharomyces Genus in Fermentation . MSystems, 6(4), 1–18. 10.1128/msystems.00260-21

Hong, K. K., & Nielsen, J. (2012). Metabolic engineering of Saccharomyces cerevisiae: A key cell factory platform for future biorefineries. Cellular and Molecular Life Sciences, 69(16), 2671–2690. 10.1007/s00018-012-0945-1

Ishchuka, O. P., Domenzain, I., Sanchez, B. J., Muñiz-Paredes, F., Martínez, J. L., Nielsen, J., & Petranovic, D. (2022). Genome-scale modeling drives 70-fold improvement of intracellular heme production in Saccharomyces cerevisiae. Proceedings of the National Academy of Sciences of the United States of America, 119(30), 1–9. 10.1073/pnas.2108245119

Jensen, N. B., Strucko, T., Kildegaard, K. R., David, F., Maury, J., Mortensen, U. H., Forster, J., Nielsen, J., & Borodina, I. (2014). EasyClone: Method for iterative chromosomal integration of multiple genes in Saccharomyces cerevisiae. FEMS Yeast Research, 14(2), 238–248. 10.1111/1567-1364.12118

Jessop-Fabre, M. M., Jakočiūnas, T., Stovicek, V., Dai, Z., Jensen, M. K., Keasling, J. D., & Borodina, I. (2016). EasyClone-MarkerFree: A vector toolkit for marker-less integration of genes into Saccharomyces cerevisiae via CRISPR-Cas9. Biotechnology Journal, 11(8), 1110–1117. 10.1002/biot.201600147

Jiao, X., Fu, X., Li, Q., Bu, J., Liu, X., Savolainen, O., Huang, L., Guo, J., Nielsen, J., & Chen, Y. (2024). De novo production of protoberberine and benzophenanthridine alkaloids through metabolic engineering of yeast. Nature Communications, 15(1), 8759. 10.1038/s41467-024-53045-3

Junker, N., Poethe, S. S., & Wendisch, V. F. (2025). Two routes for tyrosol production by metabolic engineering of Corynebacterium glutamicum. Biotechnology for Biofuels and Bioproducts, 18(1). 10.1186/s13068-025-02641-6

Karkoula, E., Melliou, E., Magiatis, P., Demopoulos, V., Karkoula, E., Magiatis, P., Melliou, E., Kotsiras, A., Mouroutoglou, C., Skantzari, A., Melliou, E., Magiatis, P., Angelis, A., Koulakiotis, N. S., Gikas, E., Halabalaki, M., Tsarbopoulos, A., Skaltsounis, A. L., Skantzari, A., … Magiatis, P. (2014). Quantitative Measurement of Major Secoiridoid Derivatives in Olive. J. Agric. Food Chem., 62(1), 600–607. http://www.ncbi.nlm.nih.gov/pubmed/23116297

Karković Marković, A., Torić, J., Barbarić, M., & Brala, C. J. (2019). Hydroxytyrosol, Tyrosol and Derivatives and Their Potential Effects on Human Health. Molecules, 24, 2001. 10.3390/molecules24102001

Koopman, F., Beekwilder, J., Crimi, B., Houwelingen, A. Van, Hall, R. D., Bosch, D., Maris, A. J. A. Van, Pronk, J. T., & Daran, J. (2012). De novo production of the flavonoid naringenin in engineered Saccharomyces cerevisiae _ Microbial Cell Factories _ Full Text. Microbial Cell Factories, 11(155), 1–15. 10.1186/1475-2859-11-155

Labun, K., Montague, T. G., Krause, M., Torres Cleuren, Y. N., Tjeldnes, H., & Valen, E. (2019). CHOPCHOP v3: Expanding the CRISPR web toolbox beyond genome editing. Nucleic Acids Research, 47(W1), W171–W174. 10.1093/nar/gkz365

Larroy, C., Fernández, M. R., González, E., Parés, X., & Biosca, J. A. (2002). Characterization of the Saccharomyces cerevisiae YMR318C (ADH6) gene product as a broad specificity NADPH-dependent alcohol dehydrogenase: relevance in aldehyde reduction. Biochemical Journal, 361(Pt 1), 163–172. http://www.ncbi.nlm.nih.gov/pmc/articles/PMC1222291/

Larroy, C., Parés, X., & Biosca, J. A. (2002). Characterization of a Saccharomyces cerevisiae NADP(H)-dependent alcohol dehydrogenase (ADHVII), a member of the cinnamyl alcohol dehydrogenase family. European Journal of Biochemistry, 269(22), 5738–5745. 10.1046/j.1432-1033.2002.03296.x

Lee, M. E., DeLoache, W. C., Cervantes, B., & Dueber, J. E. (2015). A Highly Characterized Yeast Toolkit for Modular, Multipart Assembly. ACS Synthetic Biology, 4(9), 975–986. 10.1021/sb500366v

Li, C., Hua, Y., Shi, X., Shen, W., Yang, H., Zhou, L., Xia, Y., & Chen, X. (2026). Adaptive Laboratory Evolution for Enhanced Tyrosol Tolerance Enables High-Titer Salidroside Production in Escherichia coli. ACS Synthetic Biology. 10.1021/acssynbio.6c00149

Li, J., Zhang, S., Li, H., Dai, X., Huang, C., Ma, H., Liu, H., Qi, Q., Sheng, X., & Luo, Y. (2025). Engineering phenylpyruvate decarboxylase for controlled biosynthesis of aromatic amino acid derivatives. Metabolic Engineering, 91(June), 466–479. 10.1016/j.ymben.2025.07.001

Li, X., Chen, Z., Wu, Y., Yan, Y., Sun, X., & Yuan, Q. (2018). Establishing an Artificial Pathway for Efficient Biosynthesis of Hydroxytyrosol. ACS Synthetic Biology, 7(2), 647–654. 10.1021/acssynbio.7b00385

Li, Y., Mao, J., Song, X., Wu, Y., Cai, M., Wang, H., Liu, Q., Zhang, X., Bai, Y., Xu, H., & Qiao, M. (2020). Optimization of the l-tyrosine metabolic pathway in Saccharomyces cerevisiae by analyzing p-coumaric acid production. 3 *Biotech*, *10*(6), 1–15. 10.1007/s13205-020-02223-3

Liu, Q., Yu, T., Li, X., Chen, Y., Campbell, K., Nielsen, J., & Chen, Y. (2019). Rewiring carbon metabolism in yeast for high level production of aromatic chemicals. Nature Communications, 10(1), 1–13. 10.1038/s41467-019-12961-5

Long, Y., Han, X., Meng, X., Xu, P., & Tao, F. (2024). A robust yeast chassis: comprehensive characterization of a fast-growing Saccharomyces cerevisiae. MBio, 15(2), 1–22. https://doi.org/10.1128/mbio.03196-23

López de las Hazas, M. C., Piñol, C., Macià, A., Romero, M. P., Pedret, A., Solà, R., Rubió, L., & Motilva, M. J. (2016). Differential absorption and metabolism of hydroxytyrosol and its precursors oleuropein and secoiridoids. Journal of Functional Foods, 22, 52–63. 10.1016/j.jff.2016.01.030

Lu, H., Li, F., Sánchez, B. J., Zhu, Z., Li, G., Domenzain, I., Marcišauskas, S., Anton, P. M., Lappa, D., Lieven, C., Beber, M. E., Sonnenschein, N., Kerkhoven, E. J., & Nielsen, J. (2019). A consensus S. cerevisiae metabolic model Yeast8 and its ecosystem for comprehensively probing cellular metabolism. Nature Communications, 10(1). 10.1038/s41467-019-11581-3

Luttik, M. A. H., Vuralhan, Z., Suir, E., Braus, G. H., Pronk, J. T., & Daran, J. M. (2008). Alleviation of feedback inhibition in Saccharomyces cerevisiae aromatic amino acid biosynthesis: Quantification of metabolic impact. Metabolic Engineering, 10(3–4), 141–153. 10.1016/j.ymben.2008.02.002

Maeda, H., & Dudareva, N. (2012). The shikimate pathway and aromatic amino acid biosynthesis in plants. Annual Review of Plant Biology, 63, 73–105. 10.1146/annurev-arplant-042811-105439

Mao, J., Liu, Q., Song, X., Wang, H., Feng, H., Xu, H., & Qiao, M. (2017). Combinatorial analysis of enzymatic bottlenecks of l-tyrosine pathway by p-coumaric acid production in Saccharomyces cerevisiae. Biotechnology Letters, 39(7), 977–982. 10.1007/s10529-017-2322-5

Mas, A., Guillamon, J. M., Torija, M. J., Beltran, G., Cerezo, A. B., Troncoso, A. M., & Garcia-Parrilla, M. C. (2014). Bioactive compounds derived from the yeast metabolism of aromatic amino acids during alcoholic fermentation. BioMed Research International, 2014. 10.1155/2014/898045

Milo, R. (2013). What is the total number of protein molecules per cell volume? A call to rethink some published values. BioEssays, 35(12), 1050–1055. 10.1002/bies.201300066

Minebois, R., Ramírez-Aroca, L., Balsa-Canto, E., & Querol, A. (2025). Toward Less Energy-Consuming Alcoholic Fermentations in Oenology: A Laboratory-Scale Case Study. Food Frontiers, 1–11. 10.1002/fft2.70103

Nielsen, J. (2019). Yeast Systems Biology: Model Organism and Cell Factory. Biotechnology Journal, 14(9), 1–9. 10.1002/biot.201800421

Planells-Cárcel, A., Valera-García, E., Quintas, G., Martínez, J. L., Muñiz-Calvo, S., & Guillamón, J. M. (2025). Metabolic Engineering of a Serotonin Overproducing Saccharomyces cerevisiae Strain. Microbial Biotechnology, 18(4). 10.1111/1751-7915.70140

Querol, A., Barrio, E., Huerta, T., & Ramón, D. (1992). Molecular monitoring of wine fermentations conducted by active dry yeast strains. Applied and Environmental Microbiology, 58(9), 2948–2953.

Rahmat, E., & Kang, Y. (2020). Yeast metabolic engineering for the production of pharmaceutically important secondary metabolites. Applied Microbiology and Biotechnology, 104(11), 4659–4674. 10.1007/s00253-020-10587-y

Robles-Almazan, M., Pulido-Moran, M., Moreno-Fernandez, J., Ramirez-Tortosa, C., Rodriguez-Garcia, C., Quiles, J. L., & Ramirez-Tortosa, Mc. (2018). Hydroxytyrosol: Bioavailability, toxicity, and clinical applications. Food Research International, 105(November 2017), 654–667. 10.1016/j.foodres.2017.11.053

Rodriguez, A., Kildegaard, K. R., Li, M., Borodina, I., & Nielsen, J. (2015). Establishment of a yeast platform strain for production of p-coumaric acid through metabolic engineering of aromatic amino acid biosynthesis. Metabolic Engineering, 31, 181?188-188. 10.1016/j.ymben.2015.08.003

Ruan, X., Zhang, S., Song, W., Liu, J., Chen, X., Liu, L., & Wu, J. (2022). Efficient synthesis of tyrosol from L-tyrosine via heterologous Ehrlich pathway in Escherichia coli. Chinese Journal of Chemical Engineering, 47, 18–30. 10.1016/j.cjche.2021.05.026

Rumpl, A. E. C., Goodhew, J. R., Kelly, P. F., Hirano, M., & Pyne, M. E. (2025). Brews, fuels, and opioids: Expanding the yeast Ehrlich pathway for chemical and pharmaceutical manufacturing. Biotechnology Advances, 84(May), 108684. 10.1016/j.biotechadv.2025.108684

Sánchez, B. J., Zhang, C., Nilsson, A., Lahtvee, P., Kerkhoven, E. J., & Nielsen, J. (2017). Improving the phenotype predictions of a yeast genome-scale metabolic model by incorporating enzymatic constraints. Molecular Systems Biology, 13(8), 1–16. 10.15252/msb.20167411

Sannino, F., De Martino, A., Capasso, R., & El Hadrami, I. (2013). Valorisation of organic matter in olive mill wastewaters: Recovery of highly pure hydroxytyrosol. Journal of Geochemical Exploration, 129, 34–39. 10.1016/j.gexplo.2012.10.009

Schenck, C. A., Westphal, J., Jayaraman, D., Garcia, K., Wen, J., Mysore, K. S., Ané, J. M., Sumner, L. W., & Maeda, H. A. (2020). Role of cytosolic, tyrosine-insensitive prephenate dehydrogenase in Medicago truncatula. Plant Direct, 4(5), 1–15. 10.1002/pld3.218

Scott, W. T., Smid, E. J., Notebaart, R. A., & Block, D. E. (2020). Curation and analysis of a saccharomyces cerevisiae genome-scale metabolic model for predicting production of sensory impact molecules under enological conditions. Processes, 8(9). 10.3390/PR8091195

Shen, N., Satoh, Y., Koma, D., Ohashi, H., Ogasawara, Y., & Dairi, T. (2024). Optimization of tyrosol-producing pathway with tyrosine decarboxylase and tyramine oxidase in high-tyrosine-producing Escherichia coli. Journal of Bioscience and Bioengineering, 137(2), 115–123. 10.1016/j.jbiosc.2023.12.002

Sun, J., Zhu, L., Duan, L., Li, F., Guo, S., Bian, J., & Yang, L. (2025). Metabolic engineering of Saccharomyces cerevisiae for de novo biosynthesis of hydroxytyrosol and salidroside. Applied and Environmental Microbiology, 91(8). 10.1128/aem.00712-25

Toke, D. A., Bennett, W. L., Oshiro, J., Wu, W. I., Voelker, D. R., & Carman, G. M. (1998). Isolation and characterization of the Saccharomyces cerevisiae LPP1 gene encoding a Mg2+-independent phosphatidate phosphatase. Journal of Biological Chemistry, 273(23), 14331–14338. 10.1074/jbc.273.23.14331

Tokuhiro, K., Muramatsu, M., Ohto, C., Kawaguchi, T., Obata, S., Muramoto, N., Hirai, M., Takahashi, H., Kondo, A., Sakuradani, E., & Shimizu, S. (2009). Overproduction of geranylgeraniol by metabolically engineered Saccharomyces cerevisiae. Applied and Environmental Microbiology, 75(17), 5536–5543. 10.1128/AEM.00277-09

Verho, R., Londesborough, J., Penttilä, M., & Richard, P. (2003). Engineering Redox Cofactor Regeneration for Improved Pentose Fermentation in Saccharomyces cerevisiae. Applied and Environmental Microbiology, 69(10), 5892–5897. 10.1128/AEM.69.10.5892-5897.2003

Walker, G. M., & Stewart, G. G. (2016). Saccharomyces cerevisiae in the production of fermented beverages. Beverages, 2(4), 1–12. 10.3390/beverages2040030

Wang, C., Kim, J. Y., Choi, E. S., & Kim, S. W. (2011). Microbial production of farnesol (FOH): Current states and beyond. Process Biochemistry, 46(6), 1221–1229. 10.1016/j.procbio.2011.02.020

Wang, H., Marcišauskas, S., Sánchez, B. J., Domenzain, I., Hermansson, D., Agren, R., Nielsen, J., & Kerkhoven, E. J. (2018). RAVEN 2.0: A versatile toolbox for metabolic network reconstruction and a case study on Streptomyces coelicolor. PLoS Computational Biology, 14(10), 1–17. 10.1371/journal.pcbi.1006541

Yang, C., Chen, X., Chang, J., Zhang, L., Xu, W., Shen, W., & Fan, Y. (2018). Reconstruction of tyrosol synthetic pathways in Escherichia coli. Chinese Journal of Chemical Engineering, 26(12), 2615–2621. 10.1016/j.cjche.2018.04.020

Yang, D.-D., de Billerbeck, G. M., Zhang, J.-J., Rosenzweig, F., & Francois, J.-M. (2018). Deciphering the Origin, Evolution, and Physiological Function in the Yeast Saccharomyces cerevisiae. Applied and Environmental Microbiology, 84(1), 1–16.

Zahi, M. R., Zam, W., & El Hattab, M. (2022). State of knowledge on chemical, biological and nutritional properties of olive mill wastewater. Food Chemistry, 381(January), 132238. 10.1016/j.foodchem.2022.132238

Zhang, C., & Hua, Q. (2016). Applications of genome-scale metabolic models in biotechnology and systems medicine. Frontiers in Physiology, 6(JAN), 1–8. 10.3389/fphys.2015.00413

Zhang, C., Wu, J., Sun, Q., Ding, S., Tao, H., He, Y., Qiu, H., Shu, B., Zhu, D., Zhu, H., & Hong, K. (2024). De novo production of bioactive sesterterpenoid ophiobolins in Saccharomyces cerevisiae cell factories. Microbial Cell Factories, 23(1), 1–14. 10.1186/s12934-024-02406-0

Zhang, H., Cai, P., Guo, J., Gao, J., Xie, L., Su, P., Zhai, X., Jin, B., Cui, G., Zhou, Y. J., & Huang, L. (2025). Engineering cellular dephosphorylation boosts (+)-borneol production in yeast. Acta Pharmaceutica Sinica B, 15(2), 1171–1182. 10.1016/j.apsb.2024.12.039

Zhang, J., Petersen, S. D., Radivojevic, T., Ramirez, A., Pérez-Manríquez, A., Abeliuk, E., Sánchez, B. J., Costello, Z., Chen, Y., Fero, M. J., Martin, H. G., Nielsen, J., Keasling, J. D., & Jensen, M. K. (2020). Combining mechanistic and machine learning models for predictive engineering and optimization of tryptophan metabolism. Nature Communications , 11(1). 10.1038/s41467-020-17910-1

Zwietering, M. H., Jongenburger, I., Rombouts, F. M., & Van’t Riet, K. (1990). Modeling of the bacterial growth curve. Applied and Environmental Microbiology, 56(6), 1875–1881. 10.1128/aem.56.6.1875-1881.1990

