## Supporting Information for "Designing a *Saccharomyces cerevisiae* Cell Factory for the Overaccumulation of Tyrosol using Rational Design and Genome-Scale Metabolic Modeling Approaches"

**Table S. 1. List of oligonucleotides used in this work.**

| Nº | Name | Sequence 5'→3 |
|---|---|---|
| **1** | X-2 us F | CGTCTATGAGGAGACTGTTAGTTG |
| **2** | X-2 us R | GACCACTTCGAGAGCAAGTTG |
| **3** | X-2 ds F | CCTGCATAATCGGCCTCACAG |
| **4** | X-2 ds R | CTCGCCAAGGCATTACCATC |
| **5** | CCW12p (with X-2 us) F | AACTCGCAGGCAACTTGCTCTCGAAGTGGTCCGACACGCAAAAGAAAACC |
| **6** | CCW12p R | GAGGCAAAGGATGCAAAATTTTC |
| **7** | ARO2 (with CCW12p) F | TGTGAGAAAATTTTGCATCCTTTGCCTCATGTCAACGTTTGGGAAACTGTTC |
| **8** | ARO2 R | TTAATGAACCACGGATCTGGAG |
| **9** | FBA1t (with ARO2) F | ATTTCTCCAGATCCGTGGTTCATTAAGTTAATTCAAATTAATTGATATAGTTTTTTAATG |
| **10** | FBA1t R (with X-2 ds) R | GATCCCTCTGTGAGGCCGATTATGCAGGGTAAGCTACTATGAAAGACTTTAC |
| **11** | CCW12p end F | GAGAAAATTTTGCATCCTTTGCCTC |
| **12** | FBA1t R | GTAAGCTACTATGAAAGACTTTACAAAGA |
| **13** | GV1R(ARO10) | CGTGCGAUCTATTTTTTATTTCTTTTAAGTGCCGCTGCTT |
| **14** | PG1R(TEF1p) def | ACCTGCACUTTTGTAATTAAAACTTAGATTAGATTGCTAT |
| **15** | GP2F (ARO4) | ATCTGTCAUAAAACAATGAGTGAATCTCCAATGTTCG |
| **16** | GP1F (ARO10) | AGTGCAGGUAAAACAATGGCACCTGTTACAATTGAAAAGTTC |
| **17** | PG2R (PGK1p) | ATGACAGAUTTGTTTTATATTTGTTGTAAAAAGTAGATAATT |
| **18** | GV2R (ARO4) | CACGCGAUCTATTTCTTGTTAACTTCTCTTCTTTGTCT |
| **19** | XI-3 us F | AGTTACTTGCTCTATGCGTTTGCGC |
| **20** | ADH1t (with XI-3 us) R | GCTTTCTCAGGTATAGCATGAGGTCGCTCAATCAGACGCACGCTTGGCGTCAA |
| **21** | ADH1t end F | GAGCGACCTCATGCTATACC |
| **22** | CYC1 t end R | CTTCGAGCGTCCCAAAACCT |
| **23** | CYC1t (with XI-3 ds) F | GCTTGAGAAGGTTTTGGGACGCTCGAAGTTACGTGGATTGAGCCAGCAATAC |
| **24** | XI-3 ds R | TGAGAATCCGGACCAGCAGATAATGC |
| **25** | ARO7$^{G141S}$ R | CTAGTGGCAACAGAAGAGAAGTTATTCTTA |
| **26** | ARO7$^{G141S}$ F | TAAGAATAACTTCTCTTCTGTTGCCACTAG |
| **27** | GV2R (ARO7) | CACGCGAUTTACTCTTCCAACCTTCTTAGC |
| **28** | PP1F (GPD) | ACACACCAUAGTTTATCATTATCAATACTCGCCATTTCAAAGA |
| **29** | ARO1 target1 F | GACTTTTGTTACTAATGGTTACGAGT |
| **30** | ARO1 target1 R | AAACACTCGTAACCATTAGTAACAAA |
| **31** | PDC1p R w/ARO1 inic | CTAGAATTGGGACTTTGGCTAACTGCACCATTTTGATTGATTTGACTGTGTTATTTTGCG |
| **32** | SWF1 end F | GGCCAATCTCACAGATTTAATATAG |
| **33** | ARO1 inic R | GGACACACATTCACCATGTAATG |
| **34** | ARO1 mid F | CTAGACGAGCTGTTTGAGCAAC |
| **35** | ZWF1 target2 F | GACTttCTATACAGCCAATTGGAGAG |

| | | |
|---|---|---|
| **36** | ZWF1 target2 R | AAACCTCTCCAATTGGCTGTATAGaa |
| **37** | pPSP2 F w/NAR1 end | CGTCGAAAAGGATCTCGTCTCTGTTGGGAGCACCTGGTA ATGACCCAACATCAGATGAC |
| **38** | pPSP2 R w/ZWF1 inic | TGACGGTATTTTTTTCGAATTTGACGGGGCCTTCACTCAT TTCTTCCCGTTCCAGCACTC |
| **39** | NAR1 end F | CGCTAGAGTACGTCTTCGCG |
| **40** | ZWF1 iniciR | ATCACCTGACGCACCAAAGA |
| **41** | MAE1 target1 F | GACTTTGAGTGCTTGCAAGATCACAG |
| **42** | MAE1 target1 R | AAACCTGTGATCTTGCAAGCACTCAA |
| **43** | MAE1 ko donor F | ATACTTACTCGTATATTGTGTCCAGCTTCGGATATTTGTGC TTTTGAAACATATATATTT |
| **44** | MAE1 ko donor R | AGTATTATTTTTTTTTTAAAAATAAAGAAAAAAAATTGTACAA ATATATATGTTTCAAAAG |
| **45** | MAE1 us F | AGTTTTATTATCGTACGCGTTATTG |
| **46** | MAE1 ds R | GTATGTAGATCGCTAAGCACGATG |
| **47** | ALD5 target1 F | GACTttAAGGATATTGACTACACCAG |
| **48** | ALD5 target1 R | AAACCTGGTGTAGTCAATATCCTTaa |
| **49** | ALD5 KO donor F | AGGACTTCTTGACATTTTTAGCATATATAACAATACGATAA GTGTCTCATATGAAAAATG |
| **50** | ALD5 KO donor R | TGAATATATCCTTGTTAGATTTTATCTGTATGTCACGAGGC ATTTTTCATATGAGACACT |
| **51** | ALD5 us F | GAGAATGGCTTCAAAGAACAG |
| **52** | ALD5 ds R | CTGTTTATATATCTCTCTGGCAC |
| **53** | LPP1 target3 F | GACTTTATAAGTTGTCAGGTTTCACG |
| **54** | LPP1 target3 R | AAACCGTGAAACCTGACAACTTATAA |
| **55** | REV1p Fw (with us LPP1) | TGGTCACTGGTAAGGAAAAATTCGAAAAACTACGGGACT AAGTGTTGTTATCCGATACAA |
| **56** | REV1p Rv (with LPP1) | AATACTCCTTATGTTTCTCATCCGCCATGACAGAGATCAT CGCTGGATATGCCTAGAAAT |
| **57** | SPG3 end F | AAAAATTCGAAAAACTACGGGACTAA |
| **58** | LPP1 inic R | GTTTCTCATCCGCCATGACA |
| **59** | PHA2p(target2) F | GACTTTCCATAATTGAAGGATGAATG |
| **60** | PHA2p(target2) R | AAACCATTCATCCTTCAATTATGGAA |
| **61** | REV1p F (with PHA2p us R) | GAAAATAAAAATTTAGTAGATGAATATAATACGCAGGTGT TGTTATCCGATACAAC |
| **62** | REV1p R (with PHA2p ds F) | CTCAAAGTCTTGCTGGCCATCGCTGGATATGCCTAGAAA |
| **63** | GV1R (ADH6) | CGTGCGAUCTAGTCTGAAAATTCTTTGTCGTAG |
| **64** | ADH6 F (with HHF2p) | TAGCAAAAACAACAATCAATACAATAAAATAATGTCTTATC CTGAGAAATTTG |
| **65** | HHF2p (with TDH3p) | GAAATGGCAGTATTGATAATGATGTGGAGTGTTTGCTTGG ATTC |
| **66** | GV2R (ADH7) | CACGCGAUCTATTTATGGAATTTCTTATCATAATCG |
| **67** | PV1F (HHF2p) | CACGCGAUTGTGGAGTGTTTGCTTGGATTC |
| **68** | Sc ADH6 mid F | CATTTGGCTGCTCCACTATTATG |

| | | |
|---|---|---|
| **69** | Sc ADH6 mid R | GTGGAGCAGCCAAATGTGATGG |
| **70** | TDH3p F | TCATTATCAATACTGCCATTTC |
| **71** | TDH3p (with ADH7) R | CCTGAAATTTTTCTGGGTAAAGCATTTTGTTTGTTTATGTGTGTTTATTCG |
| **72** | ADH7 F | ATGCTTTACCCAGAAAAATTTC |
| **73** | ADH7 (with ENO2t) R | GACTAATAATTCTTAGTTAAAAGCACTCTATTTATGGAATTTCTTATCATAATC |
| **74** | ENO2t F | AGTGCTTTTAACTAAGAATTATTAG |
| **75** | Sc ADH7 mid F | CTGTGGTCCAGGTAAGAGG |
| **76** | ADH1t end F | GAGCGACCTCATGCTATACC |
| **77** | MtPDH1 inic R w/HXT7p tail | CAAAAACAAAAAGTTTTTTTAATTTTAATCAAAAAATGTCTTCATCTTCAAAATCATTG |
| **78** | HXT7p end F w/MtPDH1 tail | GATTTTGAAGATGAAGACATTTTTTGATTAAAATTAAAAAAACTTTTTG |
| **79** | HXT7p change R | CTAGAACTGAAAAAAAAAAAGTATAAATACAGACGATATATGCCAATACTT |
| **80** | HXT7p change F | AGTATTGGCATATATCGTCTGTATTTATACTTTTTTTTTTTCAGTTCTAGTATATTTTG |
| **81** | HTX7p inic R | CTCGTAGGAACAATTTCGGG |
| **82** | ADH1t F /BsmBI site | CGTCTCAtgctGAGCGACCTCATGCTATACCTGAG |
| **83** | CYC1t R /BsmBI site | CGTCTCActgaCTTCGAGCGTCCCAAAACCTTC |
| **84** | URA3 5' homology F | GCTAAATTCGAGTGAAACAC |
| **85** | URA3 3' homology R | TCTTTATATTTACATGCTAAAAATGGG |
| **86** | X-3 us F | CGAGATCTTTGTGTTCGGTTAC |
| **87** | X-3 ds R | GAGGTGGTTATTGATCACCGGAAAG |
| **88** | Chr. X-4 us F | CCAAAGCTAAGAGTCCCATTTTATTC |
| **89** | Chr. X-4 ds R | CTGGTGAGGATTTACGGTATG |
| **90** | XI-1 mid us F | GTACCGTAGCCTCCTGTATACGG |
| **91** | XI-1 mid ds R | CCTAGCCAGAATGTGCAGCAGAG |
| **92** | XI-3 US F check | GCAATACATACAGGGAGTCATGGTC |
| **93** | XI-3 us F | AGTTACTTGCTCTATGCGTTTGCGC |
| **94** | XI-3 ds R | TGAGAATCCGGACCAGCAGATAATGC |
| **95** | XII-1 mid us F | GATTTAGTAGATACAATAGCACATC |
| **96** | XII-1 mid ds R | CTTGTCCTGGTACTGCATAATG |
| **97** | GPD Pro F | CGGTAGGTATTGATTGTAATTCTG |
| **98** | ADH1-test F | GAAATTCGCTTATTTAGAAGTGTC |
| **99** | CYC1-test R | CTCCTTCCTTTTCGGTTAGAG |

**Table S. 2. List of plasmids used in this work.**

| Plasmid name | Characteristics | Integration site/ORI | Marker | Source |
|---|---|---|---|---|
| **pCfB257** | USER cloning cassette-*ADH1&CYC1* terminators | Chr X-3 | *LEU2* | (Jensen et al., 2014) |
| **pCfB258** | USER cloning cassette-*ADH1&CYC1* terminators | Chr X-4 | *HIS5* | (Jensen et al., 2014) |
| **pCfB2312** | p414-*TEF1*p-Cas9-*CYC1*t | CEN/ARS | *KanR* | (Stovicek et al., 2015) |
| **pCfB3036** | USER-MarkerFree cloning-T*ADH1*&T*CYC1* | Chr XI-1 | None | (Borodina & et.al., 2016) |
| **pCfB2909** | USER cloning- USER-MarkerFree cloning-T*ADH1*&T*CYC1* | Chr XII-5 | None | (Borodina & et.al., 2016) |
| **pCfB2904** | USER-MarkerFree cloning-T*ADH1*&T*CYC1* | Chr XI-3 | None | (Borodina & et.al., 2016) |
| **pYTK096** | Pre-Assembled *URA3* integration vector (sf*GFP* dropout) | URA locus | *URA3* | (M. E. Lee et al., 2015) |
| **pCfB2904 (*ARO4** +*ARO10*)** | (P*TEF1-ARO10*) + (P*PGK1-ARO4K229L*) | Chr XI-3 | none | This work |
| **pCfB258 *ARO7**** | P*GPD-ARO7G141S* | Chr X-4 | *HIS5* | This work |
| **pCfB257 (*ADH6* + *ADH7*)** | (P*HHF2-ADH6*) + (P*TDH3-ADH7*) | Chr X-3 | *LEU2* | This work |
| **pYTK096 *PDH1*** | (T*ADH1-ARO7G141S*-P*PGK1*) + (P*TEF1-ARO4K229L*-T*CYC1*) + (T*FBA1-MtPDH1*-P*HXT7*) | URA locus | *URA3* | This work |
| **pWS158** | Cas9 and sgRNA expression backbone (sf*GFP* dropout) | 2μ | *URA3* | (Shaw et al., 2019) |
| **pWS171** | Cas9 and sgRNA expression backbone (sf*GFP* dropout) | 2μ | *LEU2* | (Shaw et al., 2019) |
| **pWS173** | Cas9 and sgRNA expression backbone (sf*GFP* dropout) | 2μ | *KanR* | (Shaw et al., 2019) |
| **pWS174** | Cas9 and sgRNA expression backbone (sf*GFP* dropout) | 2μ | *NatR* | (Shaw et al., 2019) |
| **pWS175** | Cas9 and sgRNA expression backbone (sf*GFP* dropout) | 2μ | *HygR* | (Shaw et al., 2019) |
| **p423GPD** | P*TDH3*-MCS-T*CYC1* | 2μ | *HIS3* | (Mumberg et al., 1995) |
| **p425GPD** | P*TDH3*-MCS-T*CYC1* | 2μ | *LEU2* | (Mumberg et al., 1995) |
| **p426GPD** | P*TDH3*-MCS-T*CYC1* | 2μ | *URA3* | (Mumberg et al., 1995) |

**Table S. 3. Final level-3 (L3) gene target candidates predicted by ecFactory for TyOH production in *S. cerevisiae*, ranked by priority.**

| # | Enzymes | Name | MW | Pathway | Action | K-scores | Connectivity | EV-type | EUV-comparison | Priority |
|---|---|---|---|---|---|---|---|---|---|---|
| **1** | P25377 | *ADH7* | 39.348 | sce00010_Glycolysis / Gluconeogenesis sce00040_Pentose and glucuronate interconversions sce00561_Glycerolipid metabolism sce01110_Biosynthesis of secondary metabolites | OE | 1000 | 17 | ETC | up distinct | 1 |
| **2** | Q06408 | *ARO10* | 71.384 | sce00360_Phe metabolism | OE | 1000 | 6 | ETC | up distinct | 1 |
| **3** | P20049 | *TYR1* | 50.923 | sce00400_Phe, Tyr and Trp biosynthesis sce01110_Biosynthesis of secondary metabolites | OE | 36.284 | 5 | ETC | up distinct | 1 |
| **4** | P32178 | *ARO7* | 29.747 | | | 15.856 | 2 | ETC | up distinct | 1 |
| **5** | P28777 | *ARO2* | 40.838 | | | 13.931 | 3 | ETC | up distinct | 1 |
| **6** | P08566 | *ARO1* | 174.753 | | | 13.931 | 14 | ETC | up distinct | 1 |
| **7** | Q08548 | *ALE1* | 72.227 | sce00561_Glycerolipid metabolism sce00564_Glycerophospholipid metabolism sce01110_Biosynthesis of secondary metabolites | KD | 0.259 | 19 | BC | down distinct | 1 |
| **8** | P36013 | *MAE1* | 74.375 | sce00620_Pyruvate metabolism sce01200_Carbon metabolism | KO | 0 | 7 | UP | down distinct | 1 |
| **9** | P04806 | *HXK1* | 53.738 | sce00010_Glycolysis / Gluconeogenesis sce00051_Fructose and mannose metabolism sce00052_Galactose metabolism sce00500_Starch and sucrose metabolism sce00520_Amino sugar and nucleotide sugar metabolism sce01110_Biosynthesis of secondary metabolites sce01200_Carbon metabolism | OE | 1000 | 12 | PO | up subset | 2 |
| **10** | Q12154 | *GET3* | 39.353 | sce03060_Protein export | KD | 0.261 | 4 | BC | down subset | 2 |
| **11** | P33333 | *SLC1* | 33.887 | sce00561_Glycerolipid metabolism sce00564_Glycerophospholipid metabolism sce01110_Biosynthesis of secondary metabolites | KD | 0.228 | 28 | PO | down subset | 2 |
| **12** | Q04396 | *LPP1* | 31.586 | | | 0.208 | 35 | BC | down subset | 2 |
| **13** | Q08650 | *DGA1* | 47.711 | sce00561_Glycerolipid metabolism | KD | 0.164 | 122 | PO | down subset | 2 |
| **14** | P38221 | *CDS1* | 51.822 | sce00564_Glycerophospholipid metabolism sce01110_Biosynthesis of secondary metabolites sce04070_Phosphatidylinositol signaling system | KD | 0.274 | 34 | BC | overlapped | 3 |
| **15** | P06773 | *DCD1* | 35.646 | sce00240_Pyrimidine metabolism | KD | 0.232 | 7 | BO | overlapped | 3 |
| **16** | P22133 | *MDH2* | 40.731 | sce00020_Citrate cycle (TCA cycle) sce00270_Cys and Met metabolism sce00620_Pyruvate metabolism sce00630_Glyoxylate and dicarboxylate metabolism sce01110_Biosynthesis of secondary metabolites sce01200_Carbon metabolism | KD | 0.198 | 5 | BO | overlapped | 3 |

*Abbreviations: MW, molecular weight (kDa); OE, overexpression; KD, knockdown; KO, knockout; EUV, enzyme usage variability; ETC, essential tightly constrained; BC, biomass coupled; UP, unnecessary production; PO, production optimized; BC, biomass coupled; BO, biomass optimized; EV, enzyme variability.*

*Column description: **#**, rank of the candidate; **Enzymes**, UniProt ID; **Name**, gene name; **MW**; **Pathway**, KEGG pathway identifier; **Action**, genetic modification predicted to increase TyOH production (OE, KD, or KO); **K-scores**, gene expression scores indicating the intensity and direction of the modification; **Connectivity**, connectivity of the enzyme in the metabolic network; **EV-type**, classification of the enzyme according to its variability; **EUV-comparison**, comparison of the enzyme usage variability ranges between production and biomass-optimal state; **Priority**, priority assigned to each modification for experimental implementation (1, highest priority).*

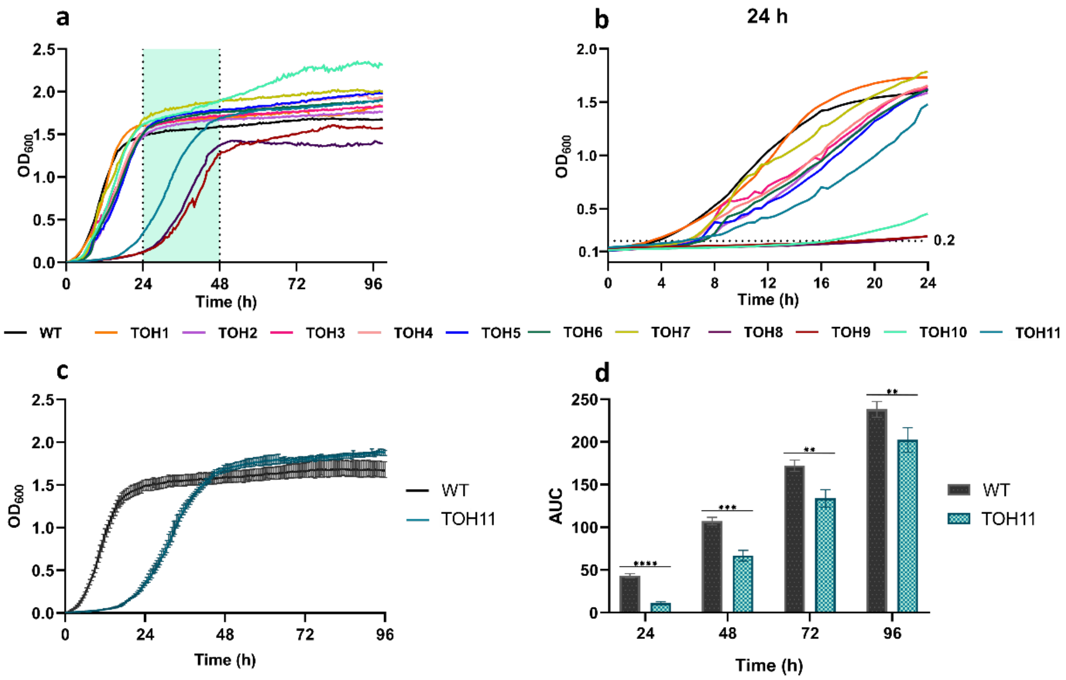


**Figure S1**. **Growth curves for each strain during the microplate experiment are represented**. **a.** The complete growth kinetics for each strain are shown, demonstrating that nearly the entire set exhibits growth comparable to that of the WT strain. **b.** The growth curve at 24 h is shown to illustrate differences in the lag phase among strains. **c**. Growth curves for the WT and TOH11 strains are plotted over time to compare their growth following cumulative modifications. **d**. AUC comparison over time between the parental strain and the TOH11 overproducer strain. Asterisks denote the significance level between WT and TOH11 at each time point, determined by two-way ANOVA with Sidak's multiple comparisons test ($p < 0.0001$ (****), $p < 0.001$ (***), $p < 0.01$ (**), $p < 0.05$ (*); ns, not significant, $p \geq 0.05$).

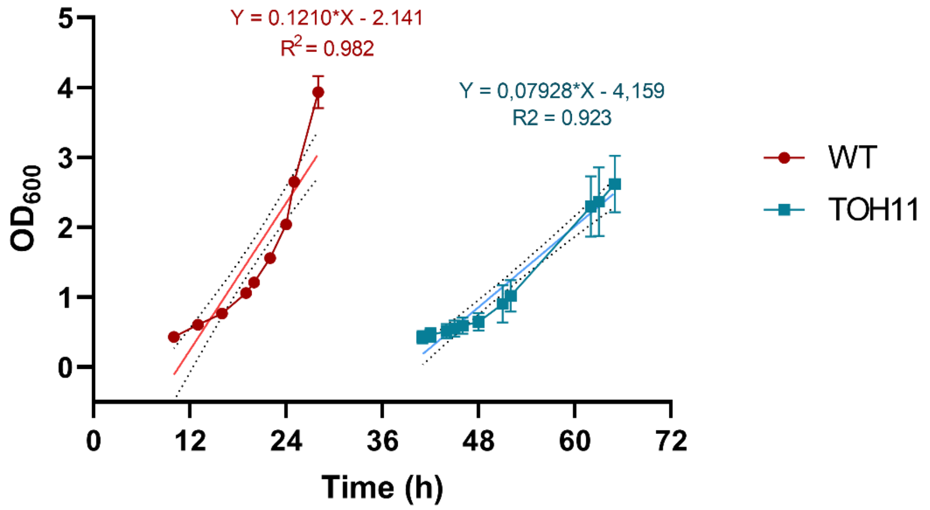


**Figure S2. Comparison of the exponential growth phase of the parental and final (TOH11) strains during batch cultivation**. Experimental $\mu_{max}$ values were determined by linear regression of ln($OD_{600}$) versus time over the exponential growth phase of each strain, with the corresponding regression equations and coefficients of determination ($R^2$) shown for each fit.

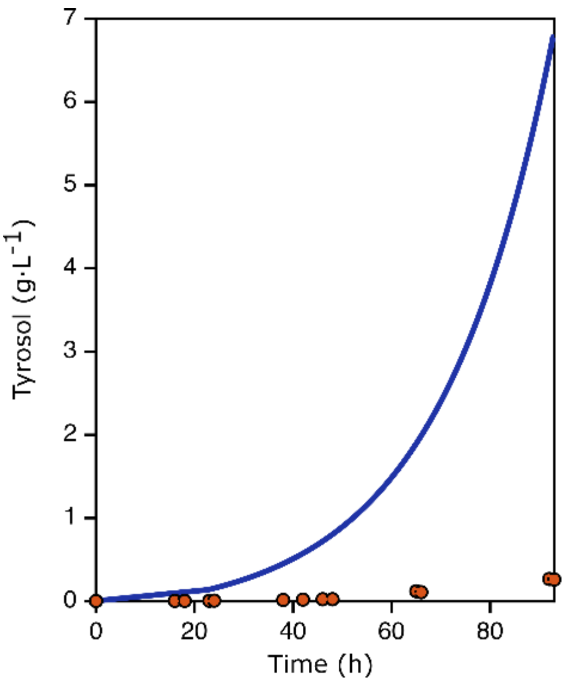


**Figure S3.** Predictive dFBA of the TOH11 batch culture relative to the ecFactory model. TyOH follows the correct increasing trend but is substantially overpredicted, consistent with unconstrained allocation of the cellular protein pool to TyOH-pathway enzymes.
